# *Drosophila* Short stop as a paradigm for the role and regulation of spectraplakins

**DOI:** 10.1101/122010

**Authors:** Andre Voelzmann, Yu-Ting Liew, Yue Qu, Ines Hahn, Cristina Melero, Natalia Sánchez-Soriano, Andreas Prokop

## Abstract

Spectraplakins are evolutionarily well conserved cytoskeletal linker molecules that are true members of three protein families: plakins, spectrins and Gas2-like proteins. Spectraplakin genes encode at least 7 characteristic functional domains which are combined in a modular fashion into multiple isoforms, and which are responsible for an enormous breadth of cellular functions. These functions are related to the regulation of actin, microtubules, intermediate filaments, intracellular organelles, cell adhesions and signalling processes during the development and maintenance of a wide variety of tissues. To gain a deeper understanding of this enormous functional diversity, invertebrate genetic model organisms, such as the fruit fly *Drosophila*, can be used to develop concepts and mechanistic paradigms that can inform the investigation in higher animals or humans. Here we provide a comprehensive overview of our current knowledge of the *Drosophila* spectraplakin Short stop (Shot). We describe its functional domains and isoforms and compare them with those of the mammalian spectraplakins dystonin and MACF1. We then summarise its roles during the development and maintenance of the nervous system, epithelia, oocytes and muscles, taking care to compare and contrast mechanistic insights across these functions in the fly, but especially also with related functions of dystonin and MACF1 in mostly mammalian contexts. We hope that this review will improve the wider appreciation of how work on *Drosophila* Shot can be used as an efficient strategy to promote the fundamental concepts and mechanisms that underpin spectraplakin functions, with important implications for biomedical research into human disease.

## 1. Introduction

The cytoskeleton comprises actin, intermediate filaments and microtubules and is essential for most, if not all, cellular processes and functions, including cell division, shape, dynamics, force generation, intracellular transport, membrane dynamics, organelle function, adhesion, signalling, cell maintenance and processes of cell death [1]. Accordingly, a high percentage of regulators of the cytoskeleton (and here we refer to components which constitute the cytoskeleton, or directly bind or associate with it) has close links to human diseases [2], and many more can be expected to be discovered in future studies.

One of the most complex and versatile protein family of cytoskeletal regulators are the spectraplakins [3, 4]. They comprise Vab10 in the worm *Caenorhabditis*, Short stop (Shot; also known as Kakapo or Groovin) in the fruit fly *Drosophila* and, in vertebrates, dystonin (also known as Bullous Pemphigoid Antigen 1/BPAG1, BP230, BP240) and Microtubule-Actin Crosslinking Factor 1 (MACF1; also known as Actin Crosslinking Family 7/ACF7, Marcrophin 1, Tabeculin α, Magellan).

Spectraplakin genes encode 7 major functional domains (Fig. 1). Through generating alternative isoforms with different combinations of these domains, spectraplakins provide a modular tool set and have been referred to as the “cytoskeleton’s Swiss army knife” [5]: they can interact with actin, intermediate filaments and microtubules alike, establish numerous structural or regulatory links between cytoskeleton components, or from cytoskeleton to other molecules or cell compartments [3, 4]. To illustrate the enormous versatility of this family, spectraplakins can be classed as true members of three important protein families:

**Fig. 1.**
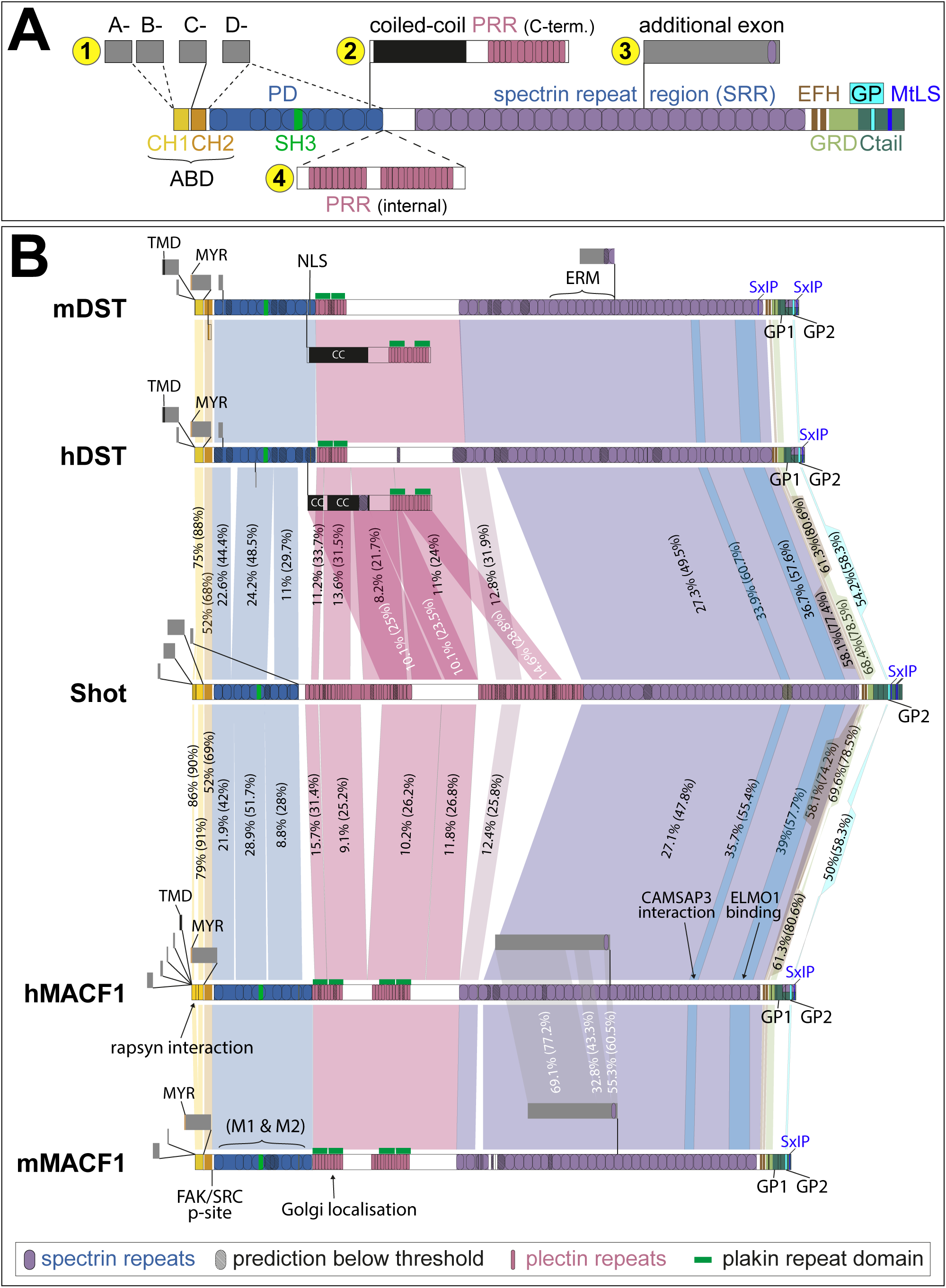
Functional domains of spectraplakins and their evolutionary conservation. **A**) Common organisation and major isoforms of spectraplakins (not to scale; abbreviations explained belox; symbols see box at bottom of B). (**1**) All spectraplakins display N-terminal variations generated through differential start sites/splicing giving rise to alternative lead sequences in combination with CH1+CH2 (A-, B-type), only CH2 (C-type) or none of both (D-type; likely also lacking the PD in Drosophila); corresponding N-termini in DST and MACF1 are referred to as type 1/2 (CH1+CH2) and type 3 (CH2 only), and isoforms lacking the entire ABD seem to be restricted to the epidermal DST isoform [3, 17]. (**2**) Short isoforms of dystonin contain a coiled-coil domain and a C-terminal PRR known to bind intermediate filaments at hemidesmosomes. (**3**) An additional exon is conserved between dystonin and MACF1 and may either be its own isoform or used as an alternative start site. (**4**) All spectraplakins display isoforms that splice in an internal PRR. **B**) Virtual full length versions of *Drosophila* Shot and human/mouse (h/m) MACF1 and dystonin (DST) displaying all known regions/domains, organised in the same way as shown in A; percentages in connecting colour beams indicate sequence identity/similarity (before/in brackets); identities/similarities between m/h versions of MACF1 and DST were not calculated; a few motifs or binding sites mentioned in the text are indicated with lines or arrows. **Abbreviations:** ABD, actin binding domain; CAMSAP, calmodulin-regulated spectrin-associated protein; CC, coiled-coil domain; CH, calponin homology domain; EFH, EF-hand domain; ELMO, engulfment and mobility; ERM, ezrin-radixin-moesin domain; FAK/Src p-site, phosphorylation site for FAK/Src; GP, GGSK-3ß phosporylation sequence; GRD, Gas2-related domain; MtLS, MT tip localisation sequence; MYR, myristoylation motif; NLS, nuclear localisation sequence; PRD, plakin repeat domain; PRR, plakin repeat region; SH3, Src homology 3 domain; SRR, spectrin repeat domain; SxIP, a MtLS consensus motif; TMD, transmembrane domain.

1. the plakins (e.g. plectin, desmoplakin, envoplakin, periplakin, epiplakin) which are cytoskeleton-associated scaffold proteins maintaining tissues under mechanical stress primarily at cell junctions [6];
2. the spectrins (e.g. α-/β-apectrin, α-actinin, dystrophin, utrophin) which primarily form links between proteins at the cell cortex [7, 8];
3. the Gas2-like proteins (Gas2, Gas2-like 1-3) which act as linkers between MTs, end binding (EB) proteins and F-actin, important for important for cytoskeletal dynamics in cell division and development [9-12].

Through their modular nature, spectraplakins functionally contribute in all three of these contexts, making them active members of those protein families. Accordingly, they have been discovered as players in a wide range of disorders or conditions. In humans, they include skin blistering of the epidermolysis bullosa simplex type (OMIM #615425) [13-16], hereditary sensory and autonomic neuropathy type VI (HSAN6; OMIM #614653) [17, 18, Edvardson, 2012 #6627], Parkinson’s disease [19, 20], neuro-developmental disorders [21-23], different forms of cancer [24-26] and the infection process of *Herpes* virus [27, 28]. Mouse models lacking *dystonin* functions have revealed additional defects in glial cells potentially linking to multiple sclerosis [29-31], and neuromuscular junction defects associated with intrinsic muscle weakness [32-34]. Mouse models lacking ACF7/MACF1 show early developmental aberration relating to Wnt signalling [35], aberrations of heart and gut physiology [36, 37], aberrations of the brain and of hair follicle stem cells both relating to cell migration defects [38-40], and defective axonal and dendritic growth [39, 41, 42].

To make sense of this enormous breadth of functions and their underlying mechanisms, invertebrate model organisms, in particular the worm *Caenorhabditis elegans* and the fruit fly *Drosophila melanogaster*, provide important experimental strategies capable of deciphering complex aspects of biology [43, 44]. These invertebrate model organisms have a long and important history of pioneering fundamental concepts and delivering understanding of molecular and biological functions, which can then be used as facilitating or instructive paradigms for related studies in higher organisms including humans [45, 46].

As an important prerequisite for applying this strategy, spectraplakins are evolutionarily well conserved (protein domains displaying up to 79% similarity (Fig. 1). In particular, the *Drosophila* spectraplakin Shot has been studied in a broad spectrum of biological contexts, revealing enormous variation in domain requirements and functional mechanisms. In a previous review we used Shot as an example to explain the methodological and experimental strategies available in *Drosophila* to decipher gene functions [44]. Here we will provide an overview of important understanding derived from such work and discuss whether and how it applies to related contexts in higher animals and humans.

## 2. A detailed comparison of the *shot* gene and its mammalian homologues

Currently, GENCODE (release 25, comprehensive gene annotation) and flybase.org (release FB2016_05) list 22 annotated isoforms for the *Drosophila shot* gene which includes 7 major, evolutionarily conserved functional domains or motifs, some containing smaller sub-motifs (Fig.1). Different isoforms can contain stark variations regarding the presence or absence of these domains. These domains include an actin-binding domain (ABD), a plakin domain (PD), a plakin repeat region (PRR), a spectrin repeat rod (SRR), two EF-hand motifs (EFH), a Gas2-related domain (GRD) and a Ctail. Most functional studies in *Drosophila* have been carried out with the Shot-PE and -PC isoforms (often referred to as Shot-LA/-LC) and deletion derivatives of these. These isoforms containing all domains except a PRR, and vary at the N-terminus (different lead sequences and a complete versus incomplete ABD; Fig. S1) [44].

The same release version of GENCODE lists 17/15 partial, overlapping and potential full length protein coding isoforms for human/mouse MACF1 and 18/14 protein coding isoforms for human/mouse dystonin (Fig. S1) which differ in part from the isoforms released by UniProt (MACF1 5 human [Q9UPN3], 4 mouse [Q9QXZ0]; DST 8 human [Q03001] and mouse [Q91ZU6]). GENCODE and UniProt are curated and validated databases, and the isoforms they list significantly deviate from the current literature [3, 17], but even in NCBI RefSeq (which lists many more automated predictions of isoform) the match is far from perfect (Tab.S1, Fig.S2). This clearly indicates that the catalogue of mammalian spectraplakins requires a systematic analysis to establish an agreed standard for the field and clarify existing controversies (e.g. about neuronal isoforms containing a C-terminal PRR) [3, 17]

### 2.1 The actin binding domain (ABD) and other N-terminal sequences

The ABD of Shot consist of two calponin homology domains (CH1 and CH2; light and dark yellow in Fig.1, respectively) [3, 4]. It displays up to 62/76% identity/similarity with that of mammalian spectraplakins, depending on the isoform and orthologue (Fig.1). According to current isoform annotations, there is not only variability in the number of CH domains but also the length and sequence of CH1 (Fig. S1).The two CH domains of fly and vertebrate spectraplakins display distinct F-actin binding properties, as is the case for most ABDs [47]: CH2 has little or no actin binding activity, CH1 alone can bind to actin but the affinity is much higher if both domains are present in tandem [3, 48-50].

Recently a point mutation in the *shot^2A2^* mutant allele, which displays a Val → Asp exchange in the IKL**V**NIR motif of CH1 (corresponding to Val^224^ in the Shot-PE isoform; Fig. S3), has been reported to abolish F-actin binding, thus confirming the central role of CH1 to this end [51]. Notably, the IKL**V**NIR motif is well conserved among ABD-containing proteins (Fig.S3). For all spectraplakins there are isoforms which either harbour a full ABD (A- and B-type isoforms in Shot; Fig. S1) or only a CH2 domain (C-type), and in some instances both CH domains can be missing (D-type) [44]. As shown in *Drosophila*, the spatiotemporal expression of such isoforms seems to correlate with actin-binding requirements of Shot functions in these regions [52] (Tab. 1).

**Table 1.** Overview of functional domain requirements during different roles of Shot. Columns list the different domains or motifs (compare Fig.1): CH1, first calponin homology; PL, plakin-like; SR, spectrin repeat; PRD, plakin-repeat; EF, EF-hand motifs; Ctail, C-terminal domain; MtLS, MT tip localization sequence (SxIP sites). Rows list different functions of Shot as explained in the text (Fas2, Fasciclin 2; GC, growth cone; “*”, localisation studies of Shot::GFP constructs in wildtype background): “+”, required; “+/-”, partially required; “−”, not required; “?”, potential role left unclear from experiments (e.g. domain was part of a rescue construct but its requirement was not further assessed); “n.d.”, never determined. References: (A) [52], (B) [134], (C) [48], (D) [42], (E) [185], (F) [66], (G) [135, 137], (H) [167].

|  | CH1 | PD | SR | PRD | EF | GRD | Ctail | MtLS |
| --- | --- | --- | --- | --- | --- | --- | --- | --- |
| **dendrite growth** <sup>(A)</sup> | + | - | - | +/- | + | + | n.d. | n.d. |
| **dendrite local.\*** <sup>(A)</sup> | - | - | +/- | n.d. | +/- | + | n.d. | n.d. |
| **axon growth** <sup>(A,B,D)</sup> | + | +/- | +/- | n.d. | +/- | + | + | + |
| **GC filopodia** <sup>(D)</sup> | - | - | - | - | + | ? | ? | n.d. |
| **Fas2 localisation** <sup>(A)</sup> | + | + | - | n.d. | + | + | n.d. | n.d. |
| **MT stability** <sup>(D)</sup> | - | - | - | n.d. | ? | + | +/- | - |
| **MT bundle org.** <sup>(B,D)</sup> | + | n.d. | n.d. | n.d. | - | + | + | + |
| **tracheal fusion** <sup>(E)</sup> | +/- | n.d. | n.d. | n.d. | n.d. | +/- | n.d. | n.d. |
| **tendon cells local.\*** <sup>(A,F)</sup> | - | - | +/- | n.d. | - | +/- | - | - |
| **tendon cell funct.** <sup>(A,F,G)</sup> | - | - | +/- | n.d. | - | + | + | - |
| **larval NMJ growth** <sup>(F)</sup> | + | - | - | n.d. | + | + | n.d. | n.d. |
| **dorsal closure local.** <sup>(H)</sup> |  |  |  |  |  |  |  |  |
| **dorsal closure** <sup>(H)</sup> | + | n.d. | n.d. | n.d. | - | + | - | - |

By default, ABDs take on a closed confirmation [53]. Recently, a FAK/Src phosphorylation site (SSL**Y**DVFP) adjacent to the CH2 domain of MACF1 was reported which promotes an open conformation and increases actin binding capability [54]. Notably, this site is conserved between MACF1, dystonin and Shot (Fig. S3) and must be given closer attention in future studies. In Shot, it might potentially relate to an auto-inhibitory mechanism (section 10) [55]; this auto-inhibition requires intramolecular association of the C-terminal EFH and GRD with the N-terminal ABD, reminiscent of anti-parallel α-actinin homodimers or spectrin heterodimers where N-terminal ABDs align with C-terminal EFH; since EFH are typical calcium-binding motifs this constellation could render F-actin-association of the ABD susceptible to calcium regulation [7, 8] (but see section 2.4). In general, there is an emerging view that ABDs can bind to more factors than F-actin [53]. So far, no further interactions were reported for Shot, but the ABD of MACF1 was shown to bind to tetratricopeptide repeat domains of rapsyn at the postsynaptic site of neuromuscular junctions [56].

As explained above, spectraplakins display different isoforms which may lack the CH1 or the complete ABD, and these isoform variations are accompanied by changes of N-terminal sequences distal to the ABD (grey in Fig.1). In Shot, these N-terminal variations (Figs.1, S1) originate from alternative transcription start sites at four distinct promoters [44, 52, 57, 58]. No specific functions were reported for these N-terminal lead sequences of Shot so far, but in dystonin they were shown to contain potential myristoylation or palmitoylation sites [50], sequences that promote dimerisation [59], and a transmembrane (TM) domain localising dystonin to perinuclear membranes [60] - although the latter report was disputed by others [50]. Current splice variant annotations listed in GENCODE contain a truncated version with the potential TM domain, but only UniProt annotates an N-terminal fragment coding for the potential myristoylation site.

Finally, both dystonin and MACF1 contain a stretch with two so-called mid segments (M1, M2) behind the ABD, which seem to co-incide with the PD; of these, the M1 domain has MT-binding capabilities [49, 61]. To our knowledge, N-terminal MT interaction has never been reported for *Drosophila* Shot.

### 2.2 The plakin domain (PD) and spectrin repeat rod (SRR)

Plakin domains (PD; blue in Fig.1) are formed by a stretch of typically 8-9 tandem spectrin repeats, each ∼100 amino acids long (varying between 70 and 130 amino acids), and a single SH3 motif [6]. The plakin domain of *Drosophila* Shot contains 9 spectrin repeats and a putative SH3 domain within the fifth repeat. It displays 23-24/42-45% identity/similarity with the plakin domains of mammalian spectraplakins (Fig.1). This is respectable when considering that spectrin repeats are highly variable in sequence (5-20% conservation), mainly defined by a common structural fold composed of two parallel and one antiparallel helix [8].

The PD of Shot was shown to be required for the compartmentalised localisation of the adhesion factor Fasciclin2 (Tab. 1) [52]. This vaguely raises the possibility that the PD can interact with transmembrane proteins (5c in Fig.2B), as was demonstrated for the PD of epidermal dystonin isoforms which bind to ß4-integrin and transmembrane collagen XVII [13, 62-64].

**Fig. 2.**
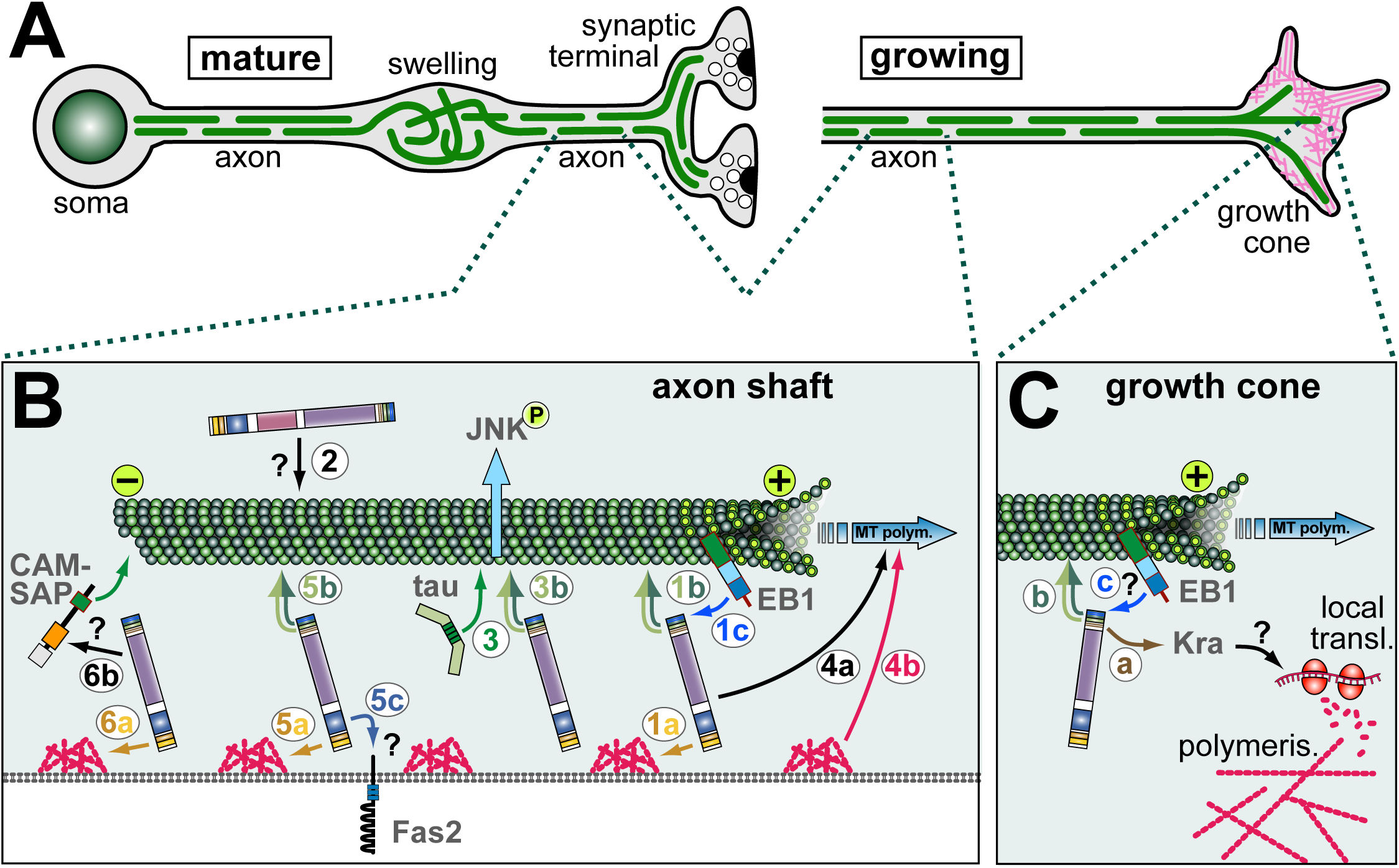
Schematic representation of known and hypothesised functions of Shot in axons. **A)** Schematics of a mature neuron (bearing a synaptic terminal; left) and a growing axon with a growth cone (right) which is rich in actin (magenta); in both cases, MTs (green lines) are arranged into parallel bundles, but in pathological swellings MTs become disorganised (e.g. during ageing or in dystonin mutant mice) [125, 126, 207]. Shot is required for the formation of these MT bundles during growth and their maintenance in mature, ageing neurons. **B)** Mechanisms of Shot in axonal MT regulation: **(1)** guidance of extending MT plus ends in parallel to the axonal surface requiring binding of ABD to cortial F-actin (1a; orange arrow), GRD/Ctail to MTs (1b; dark/light green double arrow), MtLS motifs to EB1 (1c; blue arrow); **(2)** a potential role of the PRR-containing isoform in MT bundle maintenance (directly or indirectly mediating MT bundling?); **(3)** GRD-mediated stabilisation of MTs against depolymerisation which is functionally redundant with Tau and required to maintain activated JNK in axons; **(4)** roles of Shot in promoting MT polymerisation (4a), occurring complementary to roles downstream of F-actin (4b); **(5)** roles of Shot in axonal compartmentalisation of factors (here the adhesion factor Fasciclin 2) requiring its ABD (5a), GRD (5b) and PD (5c; potentially binding the compartmentalised factor?); **(6)** extrapolating from roles in oogenesis (see Fig.3G), Shot might anchor MT minus ends linking to F-actin (6a) and CAMSAP (6b) which, in turn, binds to MT minus ends. **C)** In growth cones, Shot positively regulates the formation of F-actin rich filopodia requiring binding of C-terminal domains to MTs (b; and potentially also EB1?, c) and of its EFH to Kra (a); Kra is a putative translational regulator which may influence local actin polymerisation via local translation regulation.

Finally, the PD of dystonin contains the functional PVKRRRI/M nuclear localisation sequence which is conserved in MACF1 (PGKRRRM) [59]. Whether it is relevant in the context of the full length protein remains open. Several prediction programs identify potential nuclear localisation sequences also in the Shot PD, and N-terminal PD-containing constructs of Shot (but not full length constructs) localise to the nucleus - but this may well be an artefact caused by the GFP tag [52, 65, 66].

A second extended stretch of about 30 spectrin repeats forms the spectrin repeat rod (SRR) domain in the centre of full length Shot (purple in Fig.1), which is likewise found in both MACF1 and dystonin and is a typical element of spectrins [7, 8]. In spectrins, SRRs are elastic spacers with spring qualities, provide docking surface for protein interactions and for antiparallel dimerisation [8].

Consistent with a spacer function, the SRR of Shot is only partially required in a number of functional contexts (see Tab. 1), but indispensable for intra-molecular loop formation required for auto-inhibition (section 2.1) [55]. Dimerisation has so far not been reported for any spectraplakin, and no protein interactions have been reported for the Shot SRR. In contrast, individual spectrin repeats in the SRR of MACF1 were shown to interact with calmodulin-regulated spectrin-associated protein 3 (CAMSAP3) [67] (55% similarity with corresponding Shot repeat; Fig.S4), with ELMO [68] (58% similarity with corresponding Shot region; Fig.S4), a number of Wnt signalling pathway components (axin, LRP6, ß-catenin, GSK-3ß, APC) [35], and a region in the SRR of dystonin referred to as ERM domain was shown to bind to the dynein/Dynactin complex component p150Glued [69]. Furthermore, the SSR was shown to display functionally relevant homology to Smc family ATPases and two nucleotide-binding Walker motifs essential for the ATPase function [70]. Considering the fact that individual spectrin repeats might confer specific protein interactions, it might be functionally relevant that all fly and mammalian spectraplakins show some degree of alternative splicing (Fig. S1).

### 2.3 Plakin repeat region (PRR)

The Shot-PH isoform contains a ∼3000 amino acid PRR (pale violet in Fig. 1) between the PD and SRR which is encoded by one single exon consisting upwards of ∼40 plectin repeats (also referred to as Plakin repeats; numbers vary depending on the prediction algorithm), and a short domain stretch of 2 plectin repeats encoded by a second exon which is also contained in many other Shot isoforms [57]. Plectin repeats are typically 38 amino acids long and form a ß-hairpin followed by two alpha-helices in anti-parallel orientation [71]. In contrast to Shot (containing either 2 or ∼40 plectin repeats), the PRR of MACF1 contains ∼20, and the two separate PRRs of dystonin contain 9-11 of these motifs, respectively (8-15/21-33% identity/similarity; Fig.1). It has been proposed that modules of 4.5 plectin repeats can form a globular structure [71], or modules of five plectin repeats a solenoid structure [72]; both are referred to as plakin repeat domains (PRD) able to bind to intermediate filaments. This said, the exact features required for IF interaction remain unresolved. For example, the PRR of the *C. elegans* spectraplakin vab-10A is expected to bind intermediate filaments but is not suggested to form a PRD [6, 73].

One of the two PRR domains of dystonin localises at the C-terminus of short (hemidesmosomal) isoforms (Fig.1) and is known to link hemidesmosomes to intermediate filaments [13]. The second dystonin PRR (encoded by different exons) is localised between the PD and SRR, comparable to the PRRs of MACF1 and Shot. Little is known about the function of these internal PRRs. The first two PRDs of mouse MACF1 PRR were shown to be required for its Golgi recruitment (indicated in Fig.1B) [74]. The Shot PRR is unlikely to bind intermediate filaments, since there are no cytoplasmic intermediate filaments in arthropods (*Drosophila* has only two genes for nuclear intermediate filaments) [75, 76]. Instead, the Shot PRR localises to, and is likely required for, epidermal adherens junctions (section 11) [57], and further potential roles are suggested for the nervous system (section 4).

### 2.4 C-terminal domains: EF-hand (EFH), Gas2-related domain (GRD) and Ctail

An EF-hand is a Ca^2+^-binding motif, typically composed of a helix–loop–helix structural unit where two α-helices are bridged by a nine-residue Ca^2+^-chelation loop. In a standard EFH, the loop provides five Ca^2+^ co-ordinating groups, and two more come from a Glu or Asp located in the exiting helix (referred to as the EF-loop’s twelfth residue); further contributions are made by a hydrophobic residue following the last loop (Fig. S5) [77-82]. The EFH of dystonin and MACF1fulfill all these criteria (Fig.S5), and the EFH of dystonin was shown to mediate a strong effect on MT binding when levels of intracellular free Ca^2+^ were raised (section 13) [83]. Shot has two predicted EF-hand motifs in tandem (brown in Fig.1) which match the above structure, contain the hydrophopic residues, but provide only three co-ordinating groups in the first and five in the second EF-hand (details in Fig.S5). It is therefore difficult to predict whether the Shot EFH can bind Ca^2+^, and first functional assays (where coordination groups in the first EF-hand were mutated) were negative [55]. Instead, reported functions of the EFH of Shot comprise the mediation of protein interactions: the EF-hand mediates intra-molecular loop formation through associating with the N-terminal ABD leading to auto-inhibition (see section 2.1) [55], and it was shown to bind the putative translational regulator Krasavietz [84].

Gas2 (growth arrest specific 2)-related domains (GRD, also called GAR domains - not to be confused with the GAR domain of nucleolin; light green in Fig.1), are well conserved among spectraplakins and proteins of the Gas2-like family, and are composed of two α-helices separated by a number of extended ß-sheets [11, 66]. The GRDs of Shot, MACF1 and dystonin display up to 69/78% similarity/identity (Fig.1). All of them weakly associate with MTs in cells, but are nevertheless potent stabilisers against depolymerising drugs through still unknown mechanisms; their binding to MTs is vastly enhanced if combined with the adjacent Ctail [66, 85].

The Ctails (dark green in Fig.1) of Shot, MACF1 and dystonin also display modest association with MTs; they have no reported tertiary structure but display comparable amounts of glycins (8-10%), serins (18-24%) and arginins (11%) distributed throughout their sequences (Fig.S6); of these, the positively charged arginins were shown to be essential for MT affinity [66]. Ctails alone have no MT stabilising ability (even if their binding is strongly enhanced via a dimerisation domain) [66, 85], but they display at least two important functional features: firstly, Ctails of Shot, MACF1 and dystonin (and two of the GAS2-like proteins) contain MT tip localisation sequences (MtLS; *SxIP* or derivatives thereof; Fig.S6) which, when surrounded by positive charges, mediate binding to EB proteins and can recruit spectraplakins to the polymerising plus ends of MTs [11, 33, 66, 70, 86, 87]. Secondly, all Ctails contain a GSR repeat region (grey in Fig.S6) [85] which was later shown in MACF1 to be a functionally relevant target region for GSK-3ß phosphorylation composed of five S-R/K-X-X-S motifs (overlapping at the serines, respectively), and an additional GSK-3ß target site containing only two motifs was shown to exist further N-terminally (dark blue in Fig.S4) [38]. Similar stretches can be found in the Ctail of dystonin (two clusters) and Shot (one cluster), suggesting GSK-3ß phosphorylation to be a potential common trait (Fig.S6).

## 3. Roles of Shot during axonal pathfinding

Shot functions are best explored in the nervous system where its loss causes a variety of strong neuronal phenotypes, including defects in dendrite and axon growth, axon guidance, synapse development and maintenance, and neuronal polarity (see references below).

Axon growth reflects the ability of an axon to extend, whereas pathfinding describes the ability of a growing axon to respond to navigational cues. *In vivo*, these two very different aspects of axon development can be difficult to distinguish. For example, axon stall *in vivo* can be caused either by the general inability of the axonal growth machinery to increase axonal volume, or by the inability of the pathfinding machinery to navigate and circumvent repulsive signals. This is illustrated by analyses in *Drosophila* where actin regulator mutations (expected to affect pathfinding) cause premature stall of motoraxons in embryos, whereas the same mutations cause no phenotype or surplus axonal growth in cultured primary neurons [88].

Shot function contributes to pathfinding as well as axon growth [42, 48, 58, 66, 84, 89, 90]. Roles of Shot in pathfinding are revealed by ectopic axon crossing at the CNS midline of *shot* mutant embryos [84]. The putative translational regulator Krasavietz (Kra; homologue of human eIF-5) was shown to bind the EFH of Shot, and both proteins functionally interact during midline guidance [84]. A potential mechanism was suggested by the finding that Shot and Kra jointly promote the formation of F-actin-rich filopodia at growth cones of *Drosophila* primary neurons [42], and proper F-actin dynamics are known to be essential for axon guidance [2]. The filopodial phenotype was rescued with a C-terminal fragment of Shot comprising the EFH, GRD and Ctail (Tab. 1; Fig.1), clearly indicating that this function does not involve actin-MT linkage through Shot. The current model view is that Shot-mediated anchorage of Kra to MTs could regulate its targeted localisation to specific growth cone areas, where it can regulates local translation events known to be important during axonal pathfinding (Fig.2C) [91].

## 4. Roles of Shot during axon and dendrite growth

Healthy axons contain a core of parallel microtubule (MT) bundles (Fig.2A). However, in axons of *shot* mutant primary neurons these bundles display less coalescence, and areas of disorganised curled criss-crossing MTs are frequently observed [42]. Significant rescue of this phenotype through targeted expression of *shot* constructs only works if these constructs contain the ABD, the GRD and the Ctail with intact MtLSs (Fig.1, Tab. 1) [42, 66]. Of these, the ABD mediates binding to F-actin, likely including the cortical F-actin of axons (as is similarly the case for its close relative α-spectrin) [7, 47, 92]. GRD and Ctail are both modest MT binders, but jointly provide very strong localisation along MT shafts [66, 85]. The MtLSs interact with EB1 at polymerising plus ends of MTs, thus helping to shift a fraction of the shaft-associated Shot towards the distal MT ends [66]. We therefore proposed that these three domains establish a link between cortical F-actin and the tip of polymerising MTs, in order to guide MT extension in parallel to the axonal surface and lay MTs out into parallel bundles (1 in Fig.2B) [66].

This MT guidance model would imply that MTs are organised through Shot downstream of F-actin networks. In support of this notion, experiments where the Shot-PE isoform (containing ABD, PD, SRR, EFH, GRD and Ctail: Fig.1) was expressed in wildtype neurons, revealed a drastic increase of bundled MTs forming loops in growth cones - and this effect was suppressed upon F-destabilisation, or if the expressed construct lacked the F-actin-binding ABD domain [88]. Furthermore, when exchanging the ABD domain of Shot with other F-actin-binding domains including Lifeact [93, 94], these Lifeact-Shot hybrid proteins show unusually strong localisation all along axons (unlike endogenous Shot or transgenic Shot-PE) [42], and this phenomenon correlates with severe aberrations of MT networks (Fig.4) (Y.Q., unpublished). These findings suggest that important functions of Shot in MT regulation occur downstream of F-actin, and that the quality of its interaction with F-actin (affinity, type of actin network conformation) is important for this function.

The MT guidance model would predict that depolymerisation of cortical F-actin should lead to MT disorganisation. However, axonal MT bundles of wildtype neurons remain coalescent upon drug-induced F-actin depletion [88]. In contrast, drug-induced F-actin depletion in ‘rescued’ neurons (*shot* mutant neurons where MT bundles are rescued through expression of Shot-PE) causes severe disorganisation of axonal MTs (Y.Q. unpublished observations). The obvious difference between these two experiments is that the rescued *shot* mutant neurons express only the Shot-PE isoform (mediating MT guidance; 1 in Fig.2B), whereas the wildtype neurons harbour additional Shot isoforms that may provide resistance to F-actin loss (e.g. via MT cross-linkage; 2 in Fig.2B). A good candidate is the PRR-containing Shot-PH isoform (Fig.S1) which is strongly expressed in the nervous system including axons [44, 57].

A third potential mechanism of Shot during axon growth involves MT stabilisation against drug-induced depolymerisation via its GRD domain (section 2.4; 3 in Fig.2B). To execute its function in axons, the MT affinity of GRD is too weak and requires additional affinity provided by the adjacent Ctail (which in itself does not stabilise MTs; see section 2.4) [66]. This co-operation of GRD and Ctail appears evolutionarily conserved since it was similarly described for GRD and Ctail of mammalian spectraplakins [85].

A fourth mechanism of Shot concerns the promotion of MT polymerisation, which becomes particularly apparent if F-actin is acutely depleted in *shot* deficient neurons and MT polymerisation comes to a drastic halt (4 in Fig.2B) [95]. Shot was shown to impact on polymerisation speed through mechanisms that are independent of its ability to bind EB1 [66]. They may relate to mechanisms of other MT shaft binding proteins including Tau and MAP1B, known to impact on MT polymerisation dynamics, as was discussed previously [96].

A fifth role of Shot concerns neuronal polarisation and compartmentalisation. For example, dendritic markers were reported to be ectopically localised to axons of *shot* mutant neurons [97]. Furthermore, Fasciclin II (homologous to mammalian N-CAM) and Futsch (homologous to MAP1B), which are both known regulators of axonal growth [98-101], were shown to be aberrantly localised in neurites of *shot* mutant neurons [52, 102]. For the correct localisation of Fasciclin II, Shot requires its ABD and GRD (suggesting this to be an actin-MT linker function), but also the PD (Fig.1, Tab. 1) [52]. The PD of mammalian spectraplakins was shown to bind transmembrane proteins [13, 62, 64], and Shot might act as an anchor for certain proteins, among them membrane-associated factors (5 in Fig.2B). A recent study using different compartmentalised molecules confirmed the ABD requirement and suggested roles for Shot in organising specific MT arrangements at compartment borders within axons [103].

A sixth mechanism of Shot has recently been proposed in *Drosophila* oocytes and follicle cells and might well apply to its function in axons: Shot binds Patronin which, in turn, captures MT minus ends; since Shot can bind cortical F-actin, this mechanism anchors MTs to the plasma membrane (6 in Fig.2B; section 12) [51]. A comparable mechanism has been described for mammalian MACF1 [67, 104]. We found that *shot* mutant neurons, treated with the MT-stabilising drug taxol (promoting MT polymerisation, stabilisation and bundling) [105], display a shift of MTs to the distal end of axons (N.S.S., unpublished data; Fig.4<u>**G, H**</u>). This phenotype might be caused by | lack of minus end anchorage of MTs, in combination with axonal MT sliding performed by kinesins [106, 107].

Taken together, Shot operates in axons via a variety of mechanisms, made possible through its multi-domain structure (Fig.1). Of these molecular mechanisms, MT polymerisation and organisation (i.e. MT guidance, hypothetical MT bundle stabilisation involving PRRs, potential minus end anchorage) are likely contributors to the axonal growth-promoting roles of Shot. Also MT stabilisation against depolymerisation may contribute by helping to maintain MTs once they have been formed, thus shifting MT polymerisation towards a net plus outcome that leads to an increase in MT volume and therefore axon growth.

Apart from axon growth, also dendrite development is severely impaired upon loss of Shot function: dendrites appear stunted in all neuron classes investigated [52, 102, 108, 109]. Like in axons, Shot acts as an actin-MT linker also in dendrites (Tab. 1) [52], but the mechanistic detail remains to be unravelled. Given that dendrite growth is also stunted in MACF1 mutant mouse neurons [41], work on roles of Shot in dendrites might be of great help, especially when considering the similarities of dendrite development in flies and vertebrates [110].

## 5. Roles of Shot during axonal maintenance and synapse regulation

Some of the developmental mechanisms explained in the previous section have also been shown or suggested to contribute to roles of Shot in axon maintenance (see the “model of axon homeostasis”) [96]. For example, MT disorganisation in *shot* mutant primary neurons becomes apparent already during development and is sustained during the following days and weeks (Fig.4). Notably, this phenotype of *shot* mutant primary neurons is correlated with premature death which can be suppressed with low doses of the MT stabilising drug taxol, suggesting that MT aberrations are causative for, or at least contribute to, the cell death phenotype (N.S.S., unpublished data).

MT bundle aberration may trigger cell death through impairing axonal transport and trapping organelles. Further pathomechanisms were recently proposed to involve the Jun Kinase (JNK) pathway [111, 112]. Active JNK is found at synaptic sites in axons (Fig.2B). However, when Shot is absent, active JNK is shifted from axons to neuronal cell bodies, as was demonstrated for primary neurons in culture, as well as embryonic sensory and motor neurons [112]. The phenotype was strongly enhanced in mutant neurons lacking both Shot and Tau, another MT shaft-binding and - stabilising protein [113]. The aberrant regulation of JNK appears to be caused by defects in MT organisation and stability (referred to as MT stress), as was suggested by successful rescues of *shot-tau* mutant phenotypes when applying epothilone B, a potent MT stabilising drug [114].

Notably, the findings with JNK provide a first mechanism which can explain the strong synapse reduction previously reported for *shot* mutant neurons [102, 115]. The underlying mechanisms involve (1) the ectopic activation of JNK in neuronal cell bodies, which poses (2) a roadblock for kinesin-3 mediated axonal transport of synaptic proteins into the axon, so that kinesin-3 and its cargo become trapped in somata and synapses are starved of their supply [112]. Notably, this pathomechanisms can affect embryonic synapse formation and synapse maintenance in the adult brain alike, depending on when MT or JNK aberration is induced [112]. This function of JNK in synaptic transport is independent of its downstream transcription factor AP-1 and, instead, expected to involve direct target phosphorylation in the cytoplasm. This said, impaired Shot function has also been reported to trigger a JNK-dependent transcriptional response of AP-1 (via activation of the MAP-kinase-kinase-kinase Wallenda/DLK) which leads to surplus growth at larval neuromuscular junctions (NMJ) [111]. As explained elsewhere [116], larval NMJ growth occurs through a budding mechanism distinct from embryonic axon growth or synapse formation; this may explain how Shot-mediated signalling in different neuronal contexts can occur through AP1-independent but also AP1-dependent signalling pathways.

## 6. Comparing axonal roles of Shot and its mammalian homologues

MACF1 and dystonin are strongly expressed in the nervous system [117] where they execute essential functions during development as well as maintenance/ageing. Also Shot performs important functions during nervous system development and ageing (sections 4 and 5), and it might provide potential explanations for spectraplakin functions at both life stages. A number of neuronal phenotypes were reported in mice: (1) Loss of MACF1 impacts on neuron migration during development and (2) causes axonal projection errors *in vivo* [39-41] correlating with stalled axon growth in culture [42]. The phenotypes of dystonin deficiency comprise (3) axonal and neuronal degeneration primarily of sensory neurons, accompanied by demyelination of peripheral nerves [118], as well as (4) structural defects at neuromuscular junctions (related to muscular expression of dystonin) [56]. Importantly, both MACF1 and dystonin are strongly expressed in the brain [117] and part of their functions might be redundant, masking important aspects of each others mutant phenotypes. In contrast, dystonin expression strongly dominates in the sensory and autonomous nervous system [117], explaining the severe impact of dystonin deficiency on this part of the nervous system, as is also in agreement with human patients carrying dystonin mutations who suffer from hereditary sensory and autonomic neuropathy type VI (HSAN6; OMIM #614653) [119]. This prominent dystonin-deficient phenotype was discovered in mice as early as 1963 [120], and the extensive work performed since then has been comprehensively reviewed elsewhere [17].

Sub-cellularly, two key features were reported for degenerating *dystonin* mutant neurons. Firstly, aberrant ER and Golgi regulation is being put forward as an important pathomechanism for neurodegeneration [74, 118, 121-123], and MACF1 seems to play similar roles [124]. Secondly, a hallmark of the dystonin mutant *in vivo* phenotype is the high frequency of axon swellings displaying disorganised MTs and accumulations of organelles and intermediate filaments [118, 125, 126]. Of these, intermediate filaments were shown not to be causative for neurodegeneration [127], whereas disorganised MTs are a prominent feature observed also in primary neurons lacking dystonin [126, 128], which is clearly shared with MACF1-deficient neurons [42].

Whether axonal degeneration upon loss of dystonin is the consequence of ER/Golgi aberration, of defective MTs, or of both, needs to be resolved. Certainly, it cannot be excluded that both phenomena are interdependent, especially when considering that smooth ER extends all along axonal MTs [129, 130]. As one example for potential systemic links, dystonin loss was reported to reduce levels of protein (but not mRNA) of tau and MAP2 [126], which are important regulators of axonal MTs [113]. For *Drosophila* Shot, we have currently no insight into its potential roles during the regulation of ER/Golgi or translation in general. Instead, existing data for Shot (sections 4, 5) point at its direct roles during axonal MT regulation, including its roles during MT stabilisation (which can be rescued with C-terminal constructs localising all along axonal MTs), the direct functional interaction of Shot with EB1 in axons, the impact on MT bundle morphology when changing Shot’s interaction with F-actin, and the functional cooperation of Shot with the MT-binding protein Tau (which is known to localise primarily to axonal MTs) [110, 112, 131]. As explained in section 5, affecting these roles of Shot in MT regulation may directly contribute to neurodegeneration.

Importantly, most of the molecular mechanisms underlying the direct roles of Shot in axonal MT-regulation, have been described for mammalian spectraplakins [85, 87, 132] and provide realistic explanations for MT disorganisation and axon swellings observed in dystonin mutant mice. The human dystonin mutation which linked to type VI HSAN was reported to consist in a C-terminal frame shift deleting the GRD and Ctail of dystonin [119], hence dystonin’s ability to bind MTs - which could be relevant for roles of dystonin both in ER/Golgi and/or in axonal MT regulation alike. We propose that both possibilities should be considered in future research to improve our understanding of spectraplakins in nervous system maintenance. We would also like to point out that spectraplakins might be involved in a wider spectrum of neurodegenerative diseases, especially those linked to Tau aberration (e.g. Alzheimer’s disease, Tau-related frontotemporal dementias, dementias lacking distinctive histopathology/DLDH) [133], as deduced from Shot’s functional overlap with Tau [112].

## 7. Structural functions of Shot in specialised, force-resistant epithelial cells

Outside the nervous system, Shot function has been most intensely studied in two highly specialised epidermal cell types: (1) so called tendon cells and (2) epithelial cells in developing wings (Fig.3A,B) [52, 65, 66, 102, 134-139], as will be explained in the following.

**Fig. 3.**
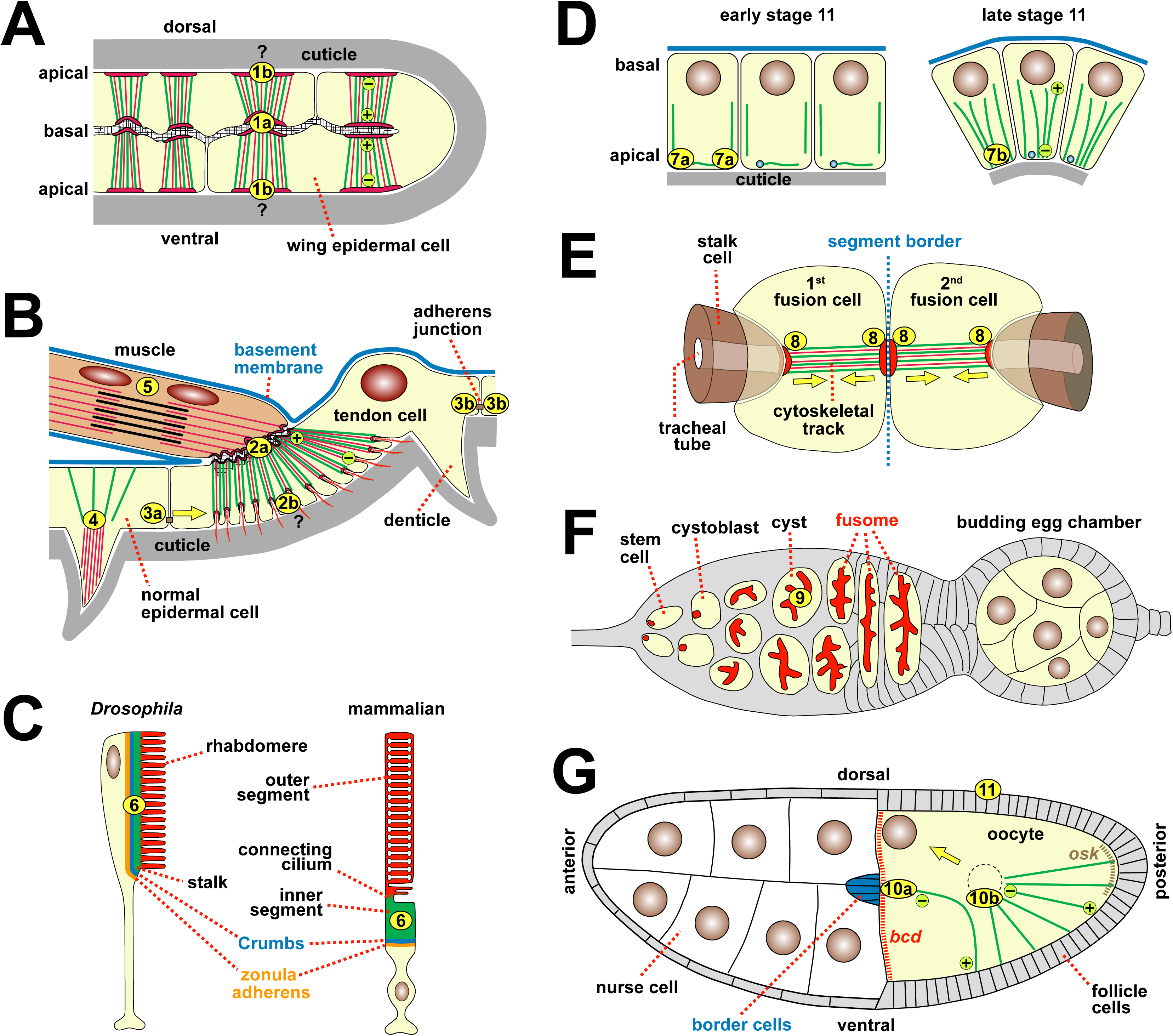
Schematic representation of cellular contexts of Shot function. **A**) The basal surfaces of developing wing epithelial cells connect via integrin-mediated cell junctions; at these junctions, Shot mediates the anchorage of apico-basal cytoskeletal arrays (1a; MTs, green; actin, magenta); it might play roles also in MT minus end anchorage (1b). **B**) Muscles attach via integrin-mediated junctions to specialised epidermal cells (tendon cells); apico-basal cytoskeletal arrays link these to cuticle on the apical surface. Shot mediates the anchorage of MT plus ends to basal junctions (2a), potentially also to apical junctions (2b); Shot plays non-autonomous roles in regulating planar polarity of epidermal denticles (3a; yellow arrow) and cell-autonomous roles in supporting denticle structure (4); it localises to (and likely stabilises) adherens junctions (3b); together with Nesprin it generates a protective MT coating around muscle nuclei (5). **C**) Photoreceptor anatomy displays evolutionarily conserved features in fly and mammals, and Shot/MACF1 localise in homologous positions (6) required for the polarity of these cells. **D**) During tube formation of salivary glands at embryonic stage 11 [208], Shot first localises from the apico-lateral (7a) to central apical (7b) positions to organise contractile actomyosin networks that reduce the apical surface (and potentially act together with CAMSAP as a MTOC (see 6 in Fig.2B). **E**) During tracheal fusion, fusion cells from two neighbouring segments (segmental border indicated by blue dashed line) meet and then pull the trailing stalk cells together (yellow arrows); Shot (8) reinforces cadherin adhesions (red) required for cytoskeletal track formation and function. **F**) During early oogenesis, Shot (9) arranges MTs at the fusome (red) required for oocyte specification. **G**) During oocyte differentiation, Shot is required for the relocation of MT minus ends and MTOC from posterior to anterior locations (10b), also required for nuclear translocation (yellow arrow) and proper localisation of polar determinants (e.g. *bicoid, bcd*, red; oskar, osk, dark green); in anterior locations, the Shot/CAMSAP complex (10a) is a key component of the MTOC; this complex plays similar roles at the apical surface of follicle cells (11). Images modified from: B [134, 149]; C) [164] D) [184]; E) [183]; F) [189].

**Fig. 4.**
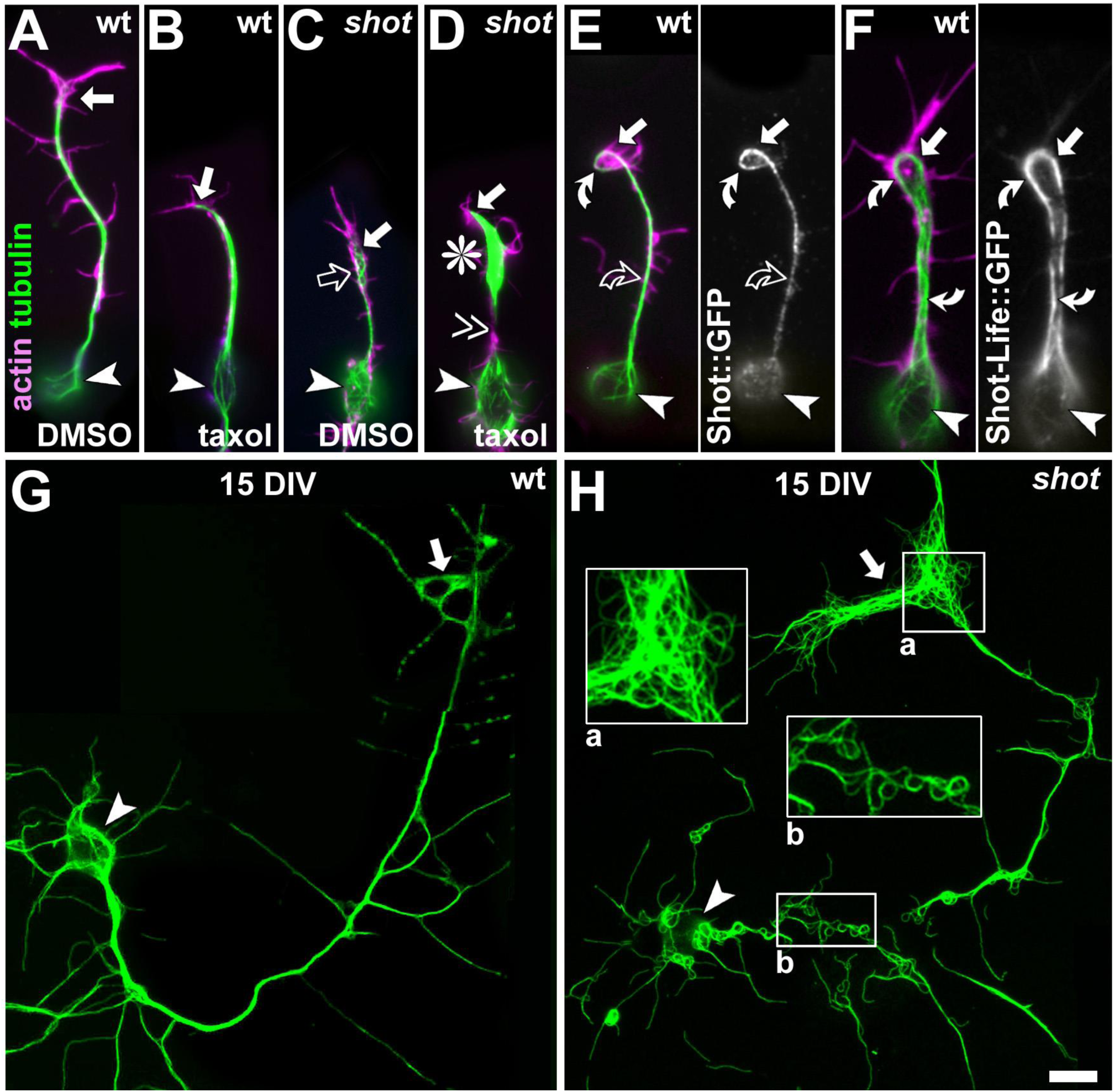
Examples of primary neurons where Shot function was manipulated. Primary neurons at 6-8 HIV (hours in vitro; A-F) or at 15 DIV (days in vitro; G,H); symbols used: white arrowheads, somata; white arrows, distal axon tips (growth cones in A-F); magenta, actin; green, tubulin. **A-D**) Wildtype (wt) neurons treated with taxol (40 nM taxol for 2 hrs; B) grow shorter than controls (treated with the vehicle DMSO; A) [88]; shot mutant control neurons (C; *shot^sf20/Df(2R)MK1^*) are shorter than wildtype and tend to display areas of MT disorganisation (open arrow) [42]; when shot mutant neurons are treated with taxol (D), tubulin staining vanishes from the axon shaft and accumulates at the distal end (asterisk). **E,F**) Wildtype neurons expressing Shot-PE::GFP show little GFP staining along the axon shaft (curved open arrow) but instead strong staining along MTs at the axon tip which have a high tendency to form bundled loops (curved white arrow) [88]; in contrast, Shot-PE^Life^::GFP localises on MTs all along axon shafts and causes not only bundled loop formation at the tip but a prominent MT bundle split all along the shaft. **G,H**) In older shot mutant neurons (H; *shot^3^*), MT disorganisation is highly abundant (close ups 2-fold magnified), whereas MTs in wildtype neurons tend to stay in bundles (G). Scale bar in H corresponds to 5 μm throughout.

The developing wing is a flat epithelial pouch (Fig.3A), the outside of which (i.e. the apical epithelial surface) is overlaid with cuticle, a specialised extracellular matrix coating the epidermal surface of most invertebrates which constitutes the exoskeleton [140]. On the inside of this flat pouch, the basal surfaces of the opposing epithelial cell layers attach to each other via integrin-mediated junctions [141]. When the wing blades unfold in the freshly hatched fly, enormous sheer and traction forces are generated. To resist these forces, wing epithelial cells contain prominent apico-basal arrays of actin filaments and MTs (red and green lines in Fig.3A) [142]. If Shot is absent from wing epithelial cells, the cells get torn apart when forces build up during wing inflation leading to blistering phenotypes in the mature wing [136, 138, 139].

Almost identical apico-basal arrays of actin filaments and MTs are found in so-called tendon cells of the epidermis which connect junctions to an apical cuticle with basal integrin-mediated junctions anchoring the tips of muscles (Fig.3B). Apart from basal integrin junctions, wing epithelial cells and tendon cells have in common that their MTs are composed of 15 protofilament (rather than the usual 13), that their MTs are nucleated at the apical cell surface, whereas their extending plus ends are captured at the basal surface [143-146], and that Shot localises all along the cytoskeletal arrays but is enriched at the basal and apical ends [52, 65, 66, 134, 135, 137, 139]. The role of Shot is expected to be the same in both cell types, but has been investigated in greater detail in tendon cells, as will be briefly summarised below.

Tendon cells require the prominent apico-basal cytoskeleton arrays as the mechanical link that transmits the pulling forces of muscles to the cuticle (Fig.3B) [134, 147, 148]. In the absence of Shot, the formation and anchorage of the basal MT arrays is defective, so that tendon cells fail to resist muscle contractions and their basal junctions get torn away [52, 65, 102, 134]. Shot contributes through at least two very distinct mechanisms: firstly, the up-regulation of tubulin expression and, secondly, the capture/anchorage of MTs.

First, during embryonic development, tubulin expression in tendon cells occurs at low levels and is severely up-regulated when myotubes approach the tendon cells [137]. Tubulin up-regulation requires EGF-receptor signalling induced by the neuregulin-like factor Vein which is secreted by the approaching myoblasts and then accumulates at the myo-epidermal junction. In *shot* mutant embryos, Vein fails to accumulate at the junction and the up-regulation of tubulin is impaired; this would explain low densities of MTs observed upon ultrastructural analysis of *shot* mutant tendon cells, expected to weaken their force resistance [102, 137]. The underlying molecular mechanisms of Shot in this context are unclear. They have been suggested to involve the anchorage of Vein receptors, the organisation of Vein-binding extracellular matrix at these junctions, or signalling cross-talk between tendon and neighbouring epidermal cells [137, 149].

Second, Shot deficiency affects the capture/anchorage of MT arrays at the basal tendon cell surface [102, 134]. In mutant tendon cells, the layer of electron dense material underlying the membrane at basal cell junctions is thinner, suggesting that Shot might be a constituent of this protein complex [102]. Importantly, the integrin-mediated junction with muscles is otherwise unaffected, suggesting that Shot is not required for junction formation but merely for linking the junction to MTs. *Vice versa*, Shot function is not dependent on integrins: basal MT attachments are maintained in integrin-deficient tendon cells, potentially orchestrated through factors involved in the early formation of myo-epidermal junctions before integrins take action [148, 150]. Whether Shot might also be required for apical anchorage of MTs in tendon cells has never been addressed, but would not be surprising when extrapolating from roles of MACF1 or Shot in localising MT minus ends to apical surfaces of epithelial cells [51, 104].

Structure-function analyses with domain deletion constructs have revealed that the localisation and function of Shot in tendon cells do not require the ABD, PD, EFH or MtLS but there is a strong dependency on GRD and Ctail and a partial requirement for the SRR (Fig.1; Tab. 1) [52, 66, 135, 137]. Therefore, Shot in this context is neither an actin-MT linker nor does it anchor via its PD to transmembrane proteins (as is the case for dystonin at hemidesmosomes) [13, 62, 64]. Future work should therefore focus not on structural linker functions but on potential other roles. For example, Shot could function as an MT stabiliser (through its C-terminus), as a factor involved in capturing MT plus ends at the basal surface (independent of its ability to bind actin or EB1), or as an orchestrator of MT arrays through mediation of signalling events (see section 13).

## 8. Comparing the roles of Shot and mammalian spectraplakins in force-resistant cells

At first sight, the phenotypes of *shot* mutant tendon cells seem comparable to that of hemidesmosomes in dystonin deficient basal keratinocytes: cytoskeletal anchorage at integrin-mediated junctions is lost, leading to cell rupture upon force application (in the case of dystonin: non-scarring skin blistering of the epidermolysis bullosa simplex type; OMIM #615425) [128, 151]. However, in contrast to hemidesmosomes where dystonin recruitment depends on integrins and requires its PD to link to the junctional adhesion factors [13, 62, 64], Shot function in tendon cells seem to work through fundamentally different mechanisms which do not depend on integrins nor on the PD (section 7). Notably, the *C. elegans* spectraplakin vab-10 at epidermal integrin junctions exists in two isoforms: a long one mediating anchorage of MTs and a short one linking to intermediate filaments [73, 152], suggesting that functions related to Shot and short epidermal isoforms of dystonin might co-exist in the same epidermal cells in worms.

A more likely mammalian context in which Shot might serve as a paradigm, is the inner ear where dystonin is expressed in some types of support cells (Deiter’s cells) which surround the hair cells of the organ of Corti [153]. These support cells sit between the basilar and tectorial membranes and are directly impacted by mechanical forces generated by pressure waves during hearing [154]. To withstand these forces, support cells display properties very similar to tendon or wing epithelial cells. They contain apico-basal (but also baso-apical) arrays of microtubules interspersed with F-actin bundles [155-158], and their microtubules display the unusual diameter of 15 rather than 13 protofilaments [159]. Since support cells play highly important roles in the organ of Corti [160, 161], work on spectraplakins in these cells might gain more attention in future, and *Drosophila* Shot might provide a helpful paradigm. Also MACF1 expression was reported in the organ of Corti, localising to hair cells where it is positioned between actin bundles of the hair and MTs coming from the cell body; so far, no functional data have been reported [162], but MACF1 might either act as a structural actin-MT linker or in regulating the apical hair morphology (similar to cell-autonomous roles of Shot in epidermal denticle formation; section 11; 4 in Fig. 3B).

## 9. Roles of fly and mammalian spectraplakins during photoreceptor development and polarity

Shot was shown to localise in developing photoreceptor cells [163]. As detailed elsewhere [164], the development and polarity of *Drosophila* photoreceptor cells display evolutionarily conserved traits with vertebrates (Fig.3C): both contain adherens junctions (forming the zonula adherens) which link neighbouring cells and separate the basal from the apical compartment. Apically, the adjacent Crumbs domain forms the stalk in *Drosophila* and the inner segment in vertebrates, followed by the more distal domain composed of the light-sensitive membranes (microvilli in *Drosophila*, ciliary discs in vertebrates; Fig.3C). Shot localises between adherens junctions, the Crumbs domain and stable/acetylated MTs, where it is functionally required: upon loss of Shot, stable MTs are disrupted, adherens junctions are mislocalised and the apical Crumbs domain is reduced leading to polarity defects that are mechanistically linked to Crumbs function [163].Targeted expression of Shot-PE (but not Shot-PE lacking the ABD) takes on an ectopic localisation in photoreceptor cells leading to polarity defects [163] (own unpublished data), suggesting that actin interaction of Shot is not required, or even counterproductive in this context. Together, the loss- and gain-of-function phenotypes of Shot clearly highlight Shot as a major orchestrator of photoreceptor morphology.

Also mammalian spectraplakins are expressed in photoreceptor cells. No functional data are available for dystonin, which is enriched in outer segments of rods during development and later also inner segments; it shows prominent co-localisation with the transmembrane collagen XVII, suggesting that it might be involved in extracellular matrix anchorage [165]. In contrast, the localisation of mouse MACF1 occurs apically in the nuclear area of developing photoreceptors, and it co-localises with the basal body beneath the connecting cilium in mature photoreceptors; its loss affects photoreceptor development and polarisation [166]: apical markers including Crumbs are disrupted and the docking of basal bodies and subsequent ciliogenesis fail, which are crucial for the formation of the outer photoreceptor segments [166]. These functions share an astonishing commonality with those of Shot in photoreceptors, making Shot a promising paradigm to unravel the underlying mechanisms in greater detail.

## 10. Roles of spectraplakins in cell migration

Little is known about roles of Shot during cell migration *in vivo*. Axonal pathfinding (section 3) relates to this topic, and of direct relevance is recent work on dorsal closure of the embryonic epidermis [167], which is a model for epithelial cell migration for example during wound healing [168]. In embryos lacking Shot function, dorsal closure is not abolished, but dynamics of the process are significantly changed, accompanied by a mildly aberrant morphology at the leading edge: defects in MT organisation (overshooting, buckling), a reduction in filopodia number (which requires actin-MT linker function, unlike filopodia number regulation in growth cones; section 3; Fig.2C and Tab. 1), and the occurrence of occasional filopodia that grow unusually long. Structure function analyses indicate a clear requirement for actin-MT linkage with key emphasis on the ABD and GRD, but not on Ctail or its MtLS motifs [167]. Interestingly, all constructs that rescue the mutant phenotype localise at the leading edge in a similar pattern as described for endogenous Shot. In our interpretation, this suggests a potential capture or anchorage mechanism, where Shot anchors to actin at the leading edge and links to MTs through its GRD; in agreement with this notion, a construct containing the ABD but lacking the GRD localises to the leading edge, but fails to rescue the mutant phenotypes.

Potential further mechanisms of Shot during cell migration were deduced from studies in S2 cell lines [55, 86, 169]. S2 cells are believed to be derived from blood cells, are about 10 μm in diameter, contain acentrosomal MT arrays and actin-rich lamellipodia, but they display poor integrin expression accompanied by lack of focal adhesion and stress fibres, weak extracellular matrix adhesion and low motility in the culture dish [170-172]. Shot functions in S2 cells favour its localisation to the MT plus end, mediated by binding of the Ctail’s MtLS motifs to EB1 [86, 169]. This localisation is made possible through an auto-inhibitory “closed conformation” in which the C-terminal EFH and GRD loop onto the N-terminal ABD, thus blocking the interaction of GRD with MTs. Only upon entering the cell periphery, is the GRD released to bind the MT lattice, so that Shot becomes an actin-MT linker that can regulate MT behaviours at the cell edge [55, 86].

Also mammalian spectraplakins are required for cell migration. Dystonin is required for myoblast and keratinocyte migration through mechanisms that are unclear but seem to involve adhesion control [33, 173, 174]. MACF1 is strongly expressed in epidermal and endothelial cells, where it predominantly localises to the shafts and plus ends of MTs [61, 175]. Knock-down of MACF1 in cultured endothelial cells causes disorientation of MTs (because they fail to trail along actin cables), and polarisation markers such as PKCζ, APC and GSK-3ß are not maintained at leading edges during migration [132, 175]. Also in keratinocytes, MTs trail along stress fibres, which is required to target and down-regulate focal adhesions, thus preventing over-adhesion and maintaining cellular dynamics. This function was proposed to require additional ATPase activity provided by MACF1’s SRR and two Walker A and B motifs within [70], as well as Src/FAK-dependent phosphorylation of the MACF1 N-terminus [54]. For efficient wound healing of the skin, hair follicle stem cells have to migrate to the site of wounding; this process requires MT regulation through MACF1, the activity of which is dynamically coordinated through GSK-3ß-mediated phosphorylation of its C-terminus (potentially downstream of Wnt signalling from the wound site) [38]. Similarly, roles of MACF1 during neuronal migration in the brain are regulated by GSK-3ß signalling [40].

Roles of MACF1 in cell migration were also studied in other cell types. In CHO LR73 cells, MT capture depends on binding of MACF1 to the Rac-activating complex of DOCK180 with engulfment and mobility (ELMO) [68]. In breast carcinoma cells induced to migrate via activation of the receptor tyrosine kinase ErbB2, activated ErbB2 inhibits GSK-3ß-mediated phosphorylation of APC and CLASP which, in turn, recruit MACF1 to the membrane [176]. In contrast, in the HeLa cancer cell line, MACF1 seems to act upstream of CLASP2 to regulate its cortical localisation [177].

Taken together, spectraplakins seem to act through a wide range of mechanisms during cell migration likely to be cell type- and context-specific. Apart from the fundamental functional domains, also various molecular features required during migration seem well conserved in Shot (FAK- and GSK-3ß target sites, ELMO binding site; Figs.S3, S4, S6). Shot already provides a useful paradigm for the mechanisms involving EB1-dependent MT guidance (1 in Fig.2B; section 4), but its full potential in the context of cell migration has by far not been reached; its studies can be easily extended to powerful *in vivo* models (e.g. border cell, germ cell or hemocyte migration) [178-180] or to improved S2 cell systems displaying true motile properties [170].

## 11. Ectodermal functions of Shot in regulating epithelia and tubulogenesis

Shot isoforms containing the PRR localise to lateral adherens junctions in the epidermis and, unlike several other isoforms, are not enriched in tendon cells; the N-terminal part of the PRR alone is sufficient to mediate this adherens junction localisation (3b in Fig.3) [57]. Correlating with this finding, severe *shot* mutant alleles which are expected to abolish the PRR-containing isoforms, cause occasional tears in the epidermis of embryos [135] and, during oogenesis, disintegration of the follicle epithelium (Fig.3G) in form of double-layering and mislocalisation of junctional proteins [57]. Whether and how the PRR contributes functionally in this context remains to be resolved. These roles of Shot in the embryonic epithelium might help to understand roles of MACF1 in regulating the columnar epithelial cell arrangements of the intestinal mucosa in mice [37] or of E-cadherin-mediated junctions of mouse keratinocytes in culture [61].

In late embryogenesis, some ventral epidermal cells of *Drosophila* produce actin-based protrusions, called denticles, on their apical surface; these denticles display position-specific differences in shape, size and orientation determined by the differential activation of the signalling pathways that orchestrate the segmentation of the embryo [150, 181]. Asymmetric localisation of Shot to apico-posterior cell contacts regulates the planar orientation of denticles in adjacent tendon cells through non-cell autonomous mechanisms, not dependent on MTs and suggestive of roles in planar polarity (3a in Fig.3B) [149]. In addition, Shot directly localises to the actin rich bundles of denticles where it acts cell-autonomously, potentially anchoring MTs to these structures (4 in Fig.3B) [149]. A similar localisation was described for MACF1 in hair cells of the inner ear [162], potentially hinting at related functions.

Tubulogenesis is a fundamental phenomenon of development, for example during the development of the lung, kidney or vasculature [182, 183], of which at least kidney and lung display spectraplakin expression [50]. In *Drosophila*, salivary glands and tracheae are specialised tubular epithelial derivatives of the ectoderm, and their formation requires Shot function constituting a good cellular model where to study spectraplakin roles during tubulogenesis. In *Drosophila* embryos, salivary gland tubules form from two epithelial placodes through a process of highly coordinated apical cell constriction and invagination [184]. Shot requires its GRD (but not ABD) to localise to actomyosin networks at the apical medial surface of placodal cells from where minus ends of MT bundles emanate in apico-basal direction; this localisation of Shot appears crucial for a pulsatile contractile activity that drives apical constriction and tube formation (Fig.3D) [184]. It has since been described that the MT minus end-binding protein CAMSAP localises to apical surfaces of salivary glands [51]. Extrapolating from functional studies in epithelial follicle cells (Fig.3G; which show a similar co-localisation of Shot and CAMSAP), both proteins are likely to anchor MT minus ends to apical surfaces and act as a MTOC [51] (see also 6 in Fig.2).

Tracheae are tubular invaginations of the epidermis which carry air to internal organs; during their embryonic development tracheal invaginations occur in every segment to then grow to each other fusing into one continuous tube which runs along the entire anterior-posterior axis of the animal [183]. The segmental tubular branches are guided by specialised tip cells, called fusion cells, which form an E-cadherin adhesion with the adjacent tracheal stalk cell and with the fusion cell of the neighbouring segment once contact is established (Fig.3E); these E-cadherin complexes are interconnected by a trans-cellular array of actin and MTs (referred to as track) required to pull stalk cells towards each other [183]. Shot localises to and stabilises E-cadherin required for fusion and proper track formation/maintenance (Fig.3E) [185, 186]. In this context, Shot may lack either its ABD or GRD, but not both at a time [185], as if it can perform linker function as a homo-oligomer.

Nothing seems to be known about potential roles of mammalian spectraplakins in tubulogenesis so far, but the detailed examples of Shot in different contexts are highly suggestive that such roles will be discovered, for which Shot will then provide helpful information.

## 12. Roles of Shot in further tissues

*Drosophila* oogenesis is a well-established model for a number of cellular processes including cell polarisation [187]. In female germ cells carrying loss-of-function mutations of *shot*, oogenesis fails before oocytes are specified [51, 188, 189]. This phenotype relates to the fusome, a specific intracellular organelle that is required for oocyte specification (9 and red in Fig.3F) [189, 190]. In Shot deficient oocytes, MTs fail to anchor at the fusome affecting downstream functions of this structure [189].

Also subsequent oocyte differentiation requires Shot. The mutant allele *shot^v104^* (lacking the Ctail; our unpublished data) [137] acts as an antimorph, i.e. its protein displays dominant negative effects. In *shot^V104/^*^+^ heterozygous mothers, oocyte specification is unaffected, but subsequent oocyte development is frequently aberrant [188]. During normal oocytes development, the MT minus ends relocate from the posterior to the anterior pole (establishing the MTOC at the anterior pole; Fig.3G). In *shot^V104/^*^+^ heterozygous mutant oocytes, there is a significant delay of anterior MTOC formation which frequently occurs ectopically on dorsal membrane surfaces, having secondary effects also on MT-dependent transport, affecting the anterior translocation of the oocyte nucleus, the anterior localisation of the polarity determinant *bcd*, and posterior transport of the polarity determinant *oskar* (Fig.3G) [188, 191, 192]. Comparable, though qualitatively different phenotypes were demonstrated for oocytes carrying the *shot^2A2^* mutant allele, a point mutation in the ABD affecting actin binding (Fig.S4) [51]. Interestingly, in cases of *shot^2A2^* mutant germ line clones where oocyte specification was successful, the subsequent oocyte differentation is aberrant: the MT minus ends relocate from the posterior to the anterior oocyte pole but fail to be captured at the anterior cell surface (with negative impact on *oskar* RNA transport and localisation to the posterior pole); this anterior MT capture requires anchorage of Shot to cortical F-actin and its association with CAMSAP, a protein binding MT minus ends [51]. Once, the MTOC is established, the Shot-CAMSP complex constitutes its core component: it traps MT fragments generated by the MT-severing protein Katanin, and these fragments are used as the seeds for new MT polymerisation [51]. Together the data for *shot^V104/^*^+^ and *shot^2A2^* suggest that guided extension of MT minus ends to the anterior pole requires the MT-binding C-terminus, whereas their cortical capture requires the N-terminal ABD and association with CAMSAP.

In vertebrates, comparable CAMSAP-dependent functions in MT minus end anchorage have been described for MACF1 [67, 104]. Furthermore, the zebrafish MACF1 (also called Magellan) anchors MTs to the oocyte cortex required for oocyte polarity: in the absence of MACF1 many structures are mislocalised, including the nucleus, germ plasm mRNAs, organelles and the Balbiani body (a structure composed of mitochondria, ER and Golgi which represents the first marker of asymmetry) [193].

Finally, Shot functions were also discovered in muscles: Shot localises to MTs surrounding muscle nuclei which, together with elastic Nesprin networks, form a protective shield against the enormous strain produced by muscle contraction (5 in Fig.3B) [194]. Similarly, dystonin has been reported to localise around nuclei of myotubes, but more likely through binding to F-actin and the nuclear envelop itself [60]. Further functions of dystonin in myocytes have been reported in the context of cell migration and neuromuscular junction differentiation [33, 34, 56, 195]. Therefore, the functions of Shot and its mammalian homologues reported in muscle tissues so far, show no obvious homology.

## 13. Signalling pathways up- and downstream of spectraplakins

The roles of spectraplakins upstream or downstream of signalling events are emerging surprisingly slowly. Indirect roles of Shot in JNK signalling were mentioned earlier (section 5). Furthermore, *Drosophila* Shot mediates Notch receptor localisation and/or stabilisation required for Notch signalling in posterior boundary cells of the proventriculus (a specialisation of the foregut) [196]. The underlying mechanisms are unclear, but they involve a feedback loop where Shot does not only promote Notch signalling but its own transcription is activated in response to Notch [196]. The N-terminus of hemidesmosomal dystonin interacts with Erbin which, in turn, interacts with ß4-integrin and Erb-B2, providing potential links between hemidesmosome assembly and Erb-B2 signalling [63]. The SRR of MACF1 interacts with a protein complex that includes Axin, GSK-3ß, ß-catenin and APC, thus mediating the regulation and translocation of these proteins required for Wnt signalling during the formation of the primitive streak, node, and mesoderm in early mouse development [35]. Similarly, MACF1 was shown to bind GSK-3ß in neurons [40], and to regulate the polarised localisation of PKCζ, APC and GSK-3ß in migrating endothelial cells [175].

A number of upstream signalling mechanisms have been described to regulate spectraplakin function. Binding of Ca^2+^ to the EF-hand motifs of dystonin can switch between associations of the dystonin C-terminus with either EB1 or MT shafts: the 1^st^ motif seems structurally required to promote a ‘closed confirmation’, seemingly independent of Ca^2+^ binding (mutation of the 1^st^ or both motifs locks constructs into an open, MT shaft binding state), whereas binding of Ca^2+^ to the 2^nd^ motif promotes an open confirmation associating with the MT shaft (mutations of the 2^nd^ motif locks constructs into a closed, EB1-binding state) [83]. Intriguingly, the EF-hand motifs of Shot can similarly switch between Shot localisation to EB1 or MT shafts in S2 cells, but through a very different mechanism requiring intra-molecular binding to the N-terminal ABD (sections 2.4, 10). Whether the Shot EFH can be regulated through calcium remains to be seen, whereas the EFH of MACF1 is well conserved with that of dystonin and is almost certain to be regulated through Ca^2+^ (details in Fig.S5). Furthermore, phosphorylation has been reported to regulate spectraplakin functions. Firstly, GSK-3ß-mediated phosphorylation of the MACF1 C-terminus (Fig.S6) detaches MACF1 from MTs, thus leading to loss of polarised MT extension and directed cell migration during wound healing and in the developing brain [38, 40], a molecular mechanism that is likely conserved between spectraplakins (Fig.S4). Secondly, Src/FAK-mediated phosphorylation of a well conserved tyrosine residue adjacent to the second CH domain of MACF1 (Fig. S3) is required for MT targeting to focal adhesions thereby promoting the proper migration of skin epidermal cells during wound healing in mouse [54].

## 14. Conclusions and future perspectives

Here we tried to give a comprehensive overview of our current knowledge about spectraplakin functions in flies and mammals, aiming to relate the detailed understanding of Shot to comparable contexts in mammals, and explore the potential of Shot as a paradigm for future studies. For a century, efficient and cost-effective research in *Drosophila* has been used in this way, pioneering and generating molecular and conceptual understanding of fundamental biology, that has then instructed the investigation in vertebrates/mammals and provided the mechanisms and molecules that have made rapid advance possible [46, 197]. Such pioneering research in *Drosophila* capitalises on a vast repertoire of existing genetic tools, strategies and detailed knowledge [198], and this is also true for work on Shot - as has been reviewed recently [44]. Importantly, there are now efficient means to become acquainted with the use of *Drosophila* as a model [199, 200], facilitating the establishment of fly research as a second line of investigation even in laboratories specialised on vertebrate/mammalian models.

Another model organism which can provide similarly efficient strategies for pioneering spectraplakin research is the worm *Caenorhabditis elegans* [43, 201] (this issue). Unfortunately, work on its well-conserved spectraplakin Vab-10 has not yet reached its full potential. So far, Vab-10 has been shown to play roles at specific contact points of gonadal tissues and during cell and nuclear migration of gonadal tip cells [202, 203], and as a structural component of ECM-linked, integrin-dependent anchoring complexes at muscle attachments that provide interesting paradigms for hemidesmosomes [73, 201, 204-206]. As summarised here, research on *Drosophila* Shot is far more advanced and has arguably become the molecularly best understood spectraplakin in the context of relevant *in vivo* functions in a wide variety of tissues, primarily concerning cell shape/dynamics, force resistance, development and ageing/maintenance. At the mechanistic level, these roles involve the regulation of actin, microtubules, cell adhesion, cell polarity and signalling, and in many instances we know the combinations of functional domains required. Merely by comparing and contrasting these functions of Shot with each other, we have unique opportunities to unravel the fundamental mechanisms of spectraplakins, which then provides a conceptual tool box that can be applied to the investigation of spectraplakins in any phylum, including the many cases where spectraplakins are linked to human disease (see introduction).

## Acknowledgements

We would like to thank our generous funders: AV is by a postdoctoral fellowship of the German Research council (DFG; VO 2071/1-1), Y-TL is supported by her parents and funds of FBMH, NS by a BBSRC grant (BB/M007456/1), IH, YQ and AP by BBSRC grants to AP (BB/L000717/1; BB/M007553/1).

***Drosophila* Short stop as a paradigm for the role and regulation of spectraplakins**

Andre Voelzmann^1^, Yu-Ting Liew^1^, Yue Qu^1^, Ines Hahn^1^, Cristina Melero^1^, Natalia Sánchez-Soriano^2^, Andreas Prokop^1,*^

1)The University of Manchester, Manchester Academic Health Science Centre, Faculty of Biology, Medicine and Health, School of Biology, Michael Smith Building, Oxford Road, Manchester M13 9PT UK

2)University of Liverpool, Institute of Translational Medicine, Department of Cellular and Molecular Physiology, Crown Street, Liverpool, L69 3BX, UK

## Supplementary materials

**Fig. S1:**
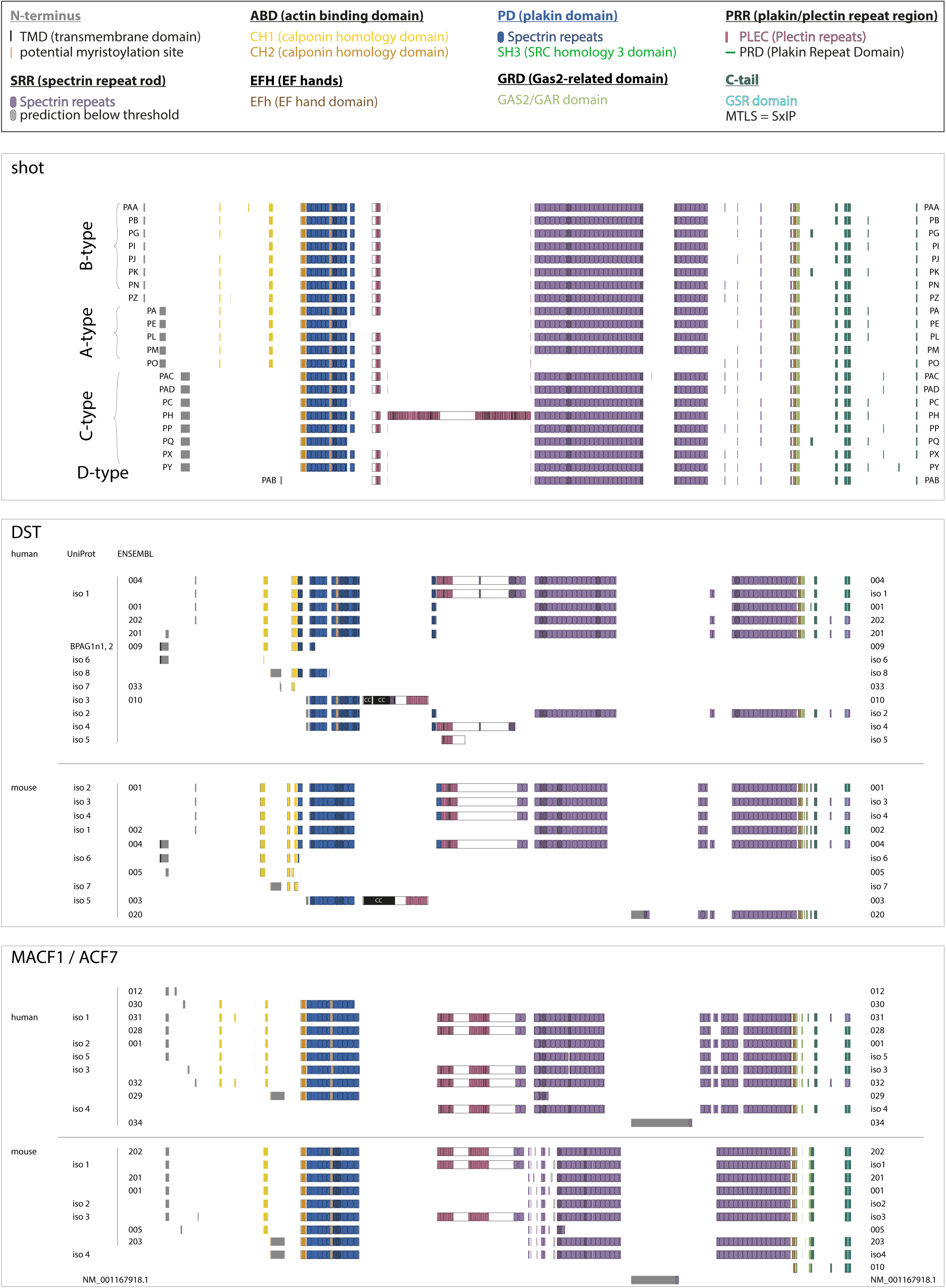
Depiction of protein modules of *shot*, DST and MACF1 isoforms currently curated and annotated in FlyBase (FB2017_01), the Comprehensive Gene Annotation datasets of GENCODE (Rel. 25) and UniProt (v10/01/2017). Isoforms completely covered by larger isoforms were omitted. Note that RefSeq GFF3 annotation predicts a much larger number of isoforms. For colour codes and symbols see Fig.1. Accession numbers used:

**Shot: *Flybase accession numbers for Drosophila melanogaster shot:*** PA = FBpp0086745, shot-PA; PB = FBpp0086742, CG18076-PB; PC = FBpp0086746, CG18076-PC; PE = FBpp0086744, CG18076-PE; PG = FBpp0086743, CG18076-PG; PH = FBpp0086747, CG18076-PH; PI = FBpp0271730, shot-PI; PJ = FBpp0271731, shot-PJ; PK = FBpp0271732, shot-PK; PL = FBpp0271733, shot-PL; PM = FBpp0271734, shot-PM; PN = FBpp0290806, shot-PN; PO = FBpp0290807, shot-PO; PP = FBpp0290808, shot-PP; PQ = FBpp0291176, CG18076-PQ; PX = FBpp0293387, shot-PX; PY = FBpp0293388, shot-PY; PZ = FBpp0293389, shot-PZ; PAA = FBpp0293390, shot-PAA; PAB = FBpp0293391, shot-PAB; PAC = FBpp0293392, shot-PAC; PAD = FBpp0293393, shot-PAD

**hMACF1: *ENSEMBL accession numbers for GENCODE 25 annotations of human MACF1:***

001 = ENST00000361689.6, MACF1-001; 012 = ENST00000484793.5, MACF1-012; 028 = ENST00000372915.7, MACF1-028; 029 = ENST00000530262.5, MACF1-029; 030 = ENST00000524432.5, MACF1-030; 031 = ENST00000567887.5, MACF1-031; 032 = ENST00000564288.5, MACF1-032; 034 = ENST00000530275.2, MACF1-034

***UniProt accession numbers for human MACF1 annotations:***

iso 1 = isoform 1, Q9UPN3-1,; iso 2 = isoform 2, Q9UPN3-2,; iso 3 = isoform 3, Q9UPN3-3; iso 4 = isoform 4, Q9UPN3-5; iso 5 = isoform 5, Q9UPN3-4

**mMACF1: *ENSEMBL accession numbers for GENCODE 25 annotations of mouse MACF1:***

001 = ENSMUST00000082108.11, Macf1-001; 005 = ENSMUST00000147030.1, Macf1-005; 201 = ENSMUST00000084301.11, Macf1-201; 202 = ENSMUST00000097897.10, Macf1-202; 203 = ENSMUST00000106220.8, Macf1-203

***RefSeq GFF3 accession number: NM_001167918.1***

***UniProt accession numbers for mouse MACF1 annotations***: iso 1 = isoform 1, Q9QXZ0-1; iso 2

= isoform 2, Q9QXZ0-2; iso = isoform 3, Q9QXZ0-3; iso 4 = isoform 4, Q9QXZ0-4

**hDST: *ENSEMBL accession numbers for GENCODE 25 annotations of human DST: 001 =***

ENST00000370788.6, DST-001; 004 = ENST00000361203.7, DST-004; 009 ENST00000449297.6, = DST-009; 010 ENST00000370765.10, = DST-010; 033 = ENST00000633795.1, DST-033; 201 = ENST00000312431.10, DST-201; 202 = ENST00000421834.6, DST-202

***UniProt accession numbers for human DST annotations:***

iso 1 = isoform 1, Q03001-7; iso 2 = isoform 2, Q03001-8; iso = isoform 3, Q03001-3; iso 4 = isoform 4, Q03001-9; iso 5 = isoform 5, Q03001-10; iso 6 = isoform 6, Q03001-11; iso 7 = isoform 7, Q03001-12; iso 8 = isoform 8, Q03001-13

**mDST: *ENSEMBL accession numbers for GENCODE 25 annotations of mouse DST***: 001 = ENSMUST00000097785.9, Dst-001; 002 = ENSMUST00000097786.9, Dst-002; 004 = ENSMUST00000183034.4, Dst-004; 005 = ENSMUST00000182697.7, Dst-005; 020 = ENSMUST00000194192.2, Dst-020

***UniProt accession numbers for human DST annotations***:

iso 1 = isoform 1, Q91ZU6-1; iso 2 = isoform 2, Q91ZU6-2; iso 3 = isoform 3, Q91ZU6-3; iso 4 = isoform 4, Q91ZU6-4; iso 5 = isoform 5, Q91ZU6-5; iso 6 = isoform 6, Q91ZU6-6; iso 7 = isoform 7, Q91ZU6-8

**Fig. S2.**
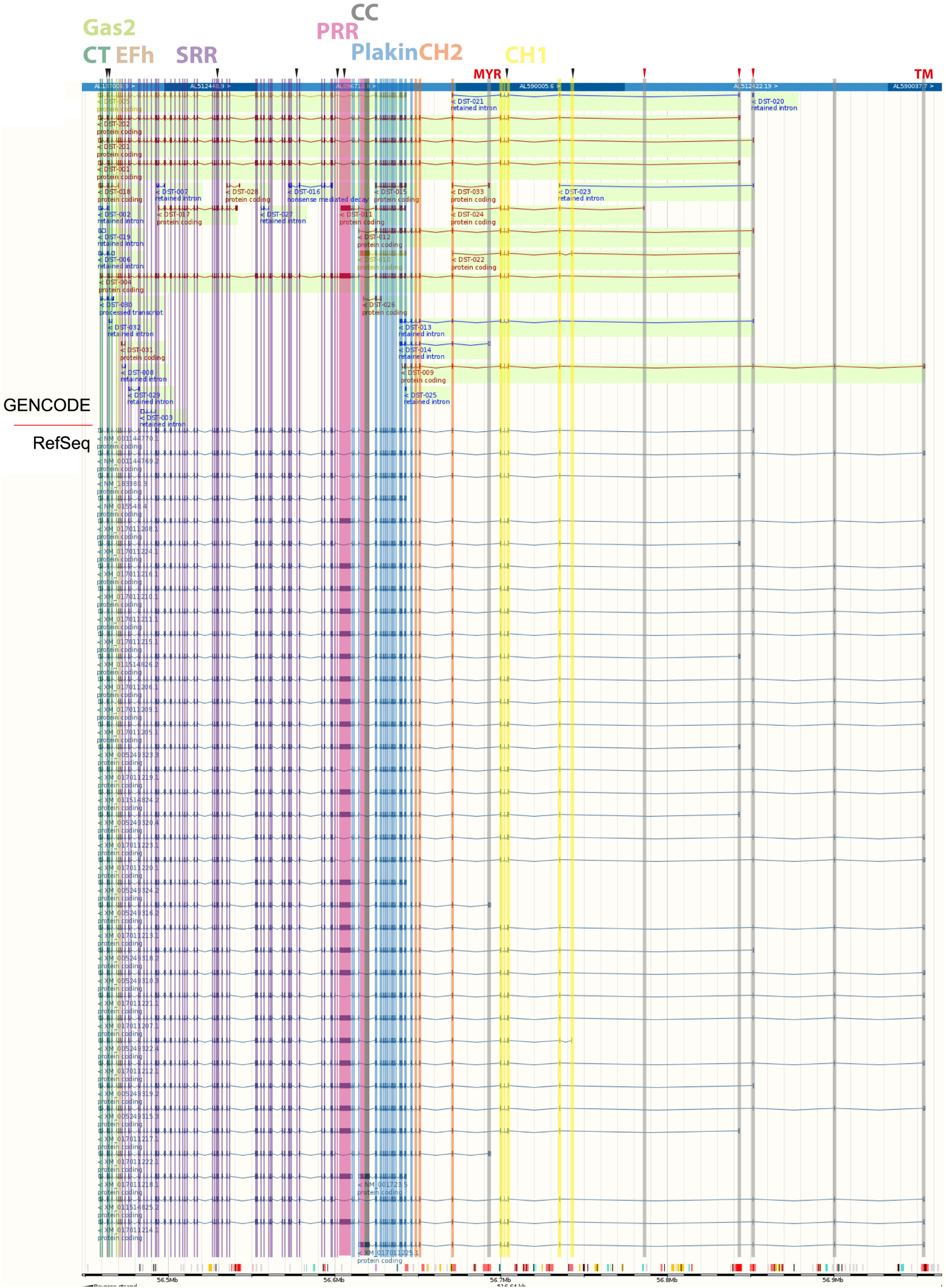
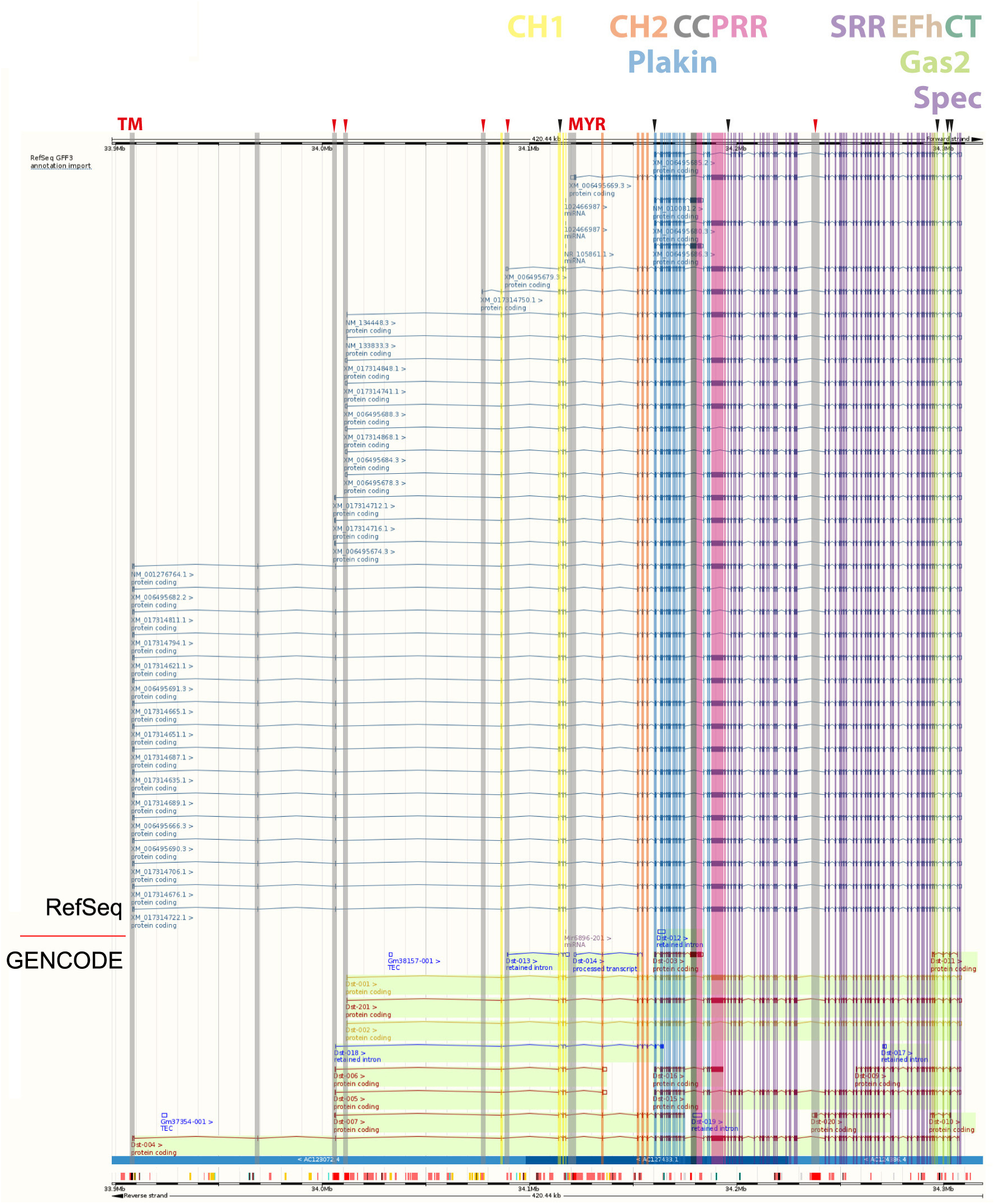
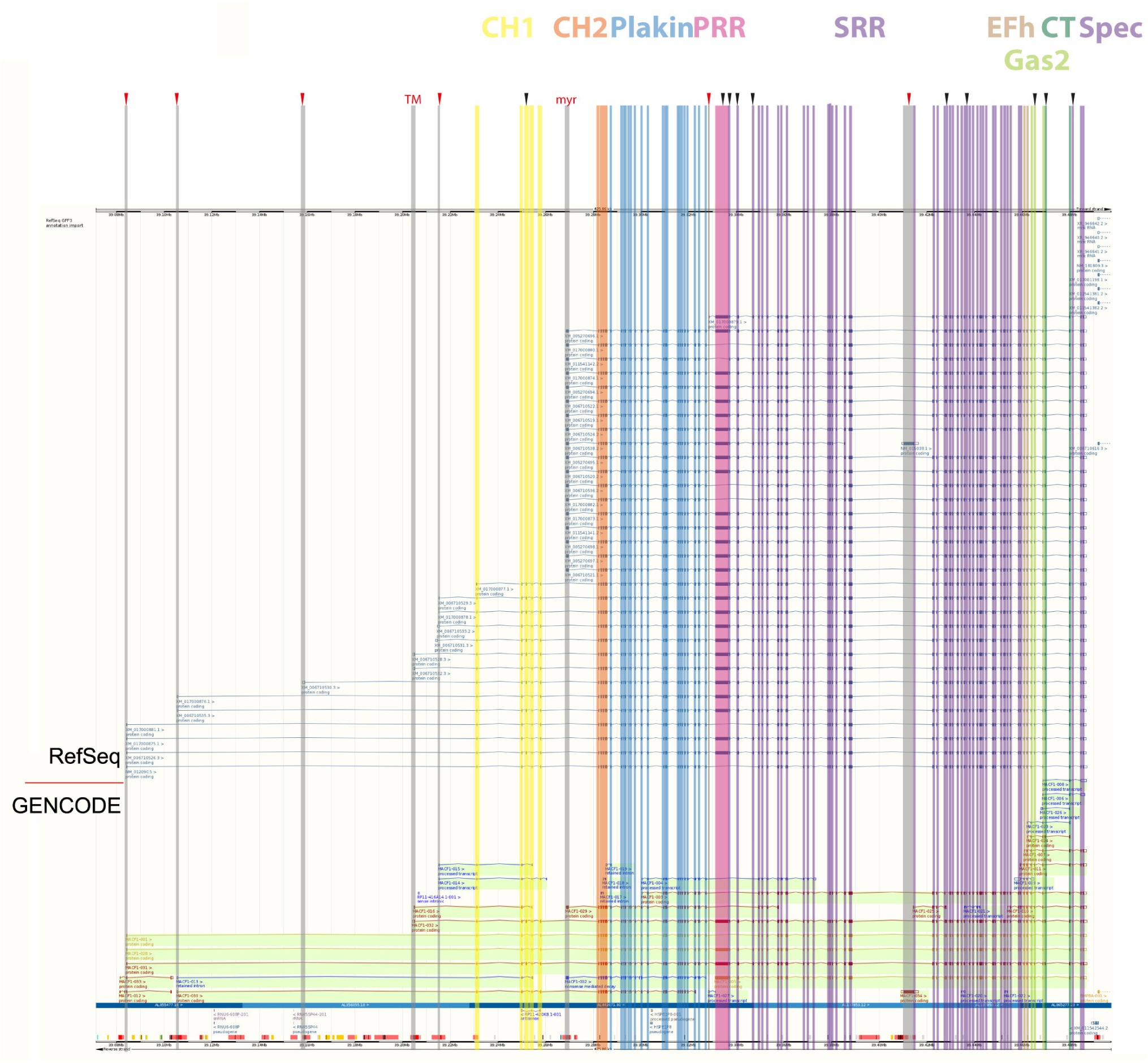
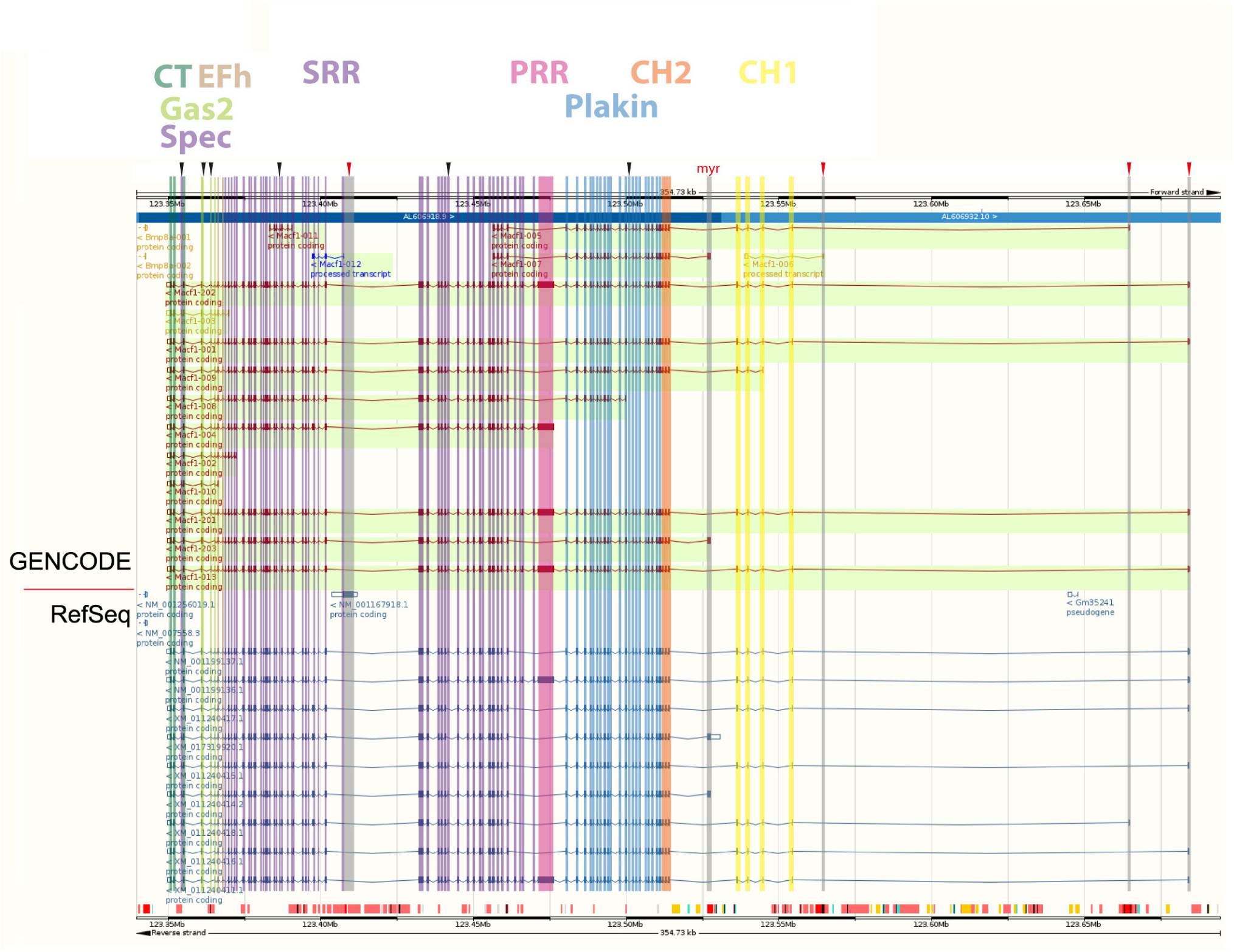
DST and MACF1 isoforms. Annotated isoforms for human/mouse (h/m) DST and MACF1 as listed on GENCODE (curated) and RefSeq (predicted; above/below red line as indicated, respectively) were obtained via the ENSEMBL genome browser (release 87) by activating the Comprehensive Gene Annotations from GENCODE 25 and RefSeq GFF3 annotation import tracks in the page configuration settings. Vertical lines match the size of coding exons and are colour-coded according to the functional domain they contribute to (same colour code as in Figs. 1 and S1). In this way, it can be easily spotted whether certain domains are present or absent in specific isoforms. Potential matches of isoforms described in the literature [1, 2] could be found on GENCODE 25 (G), UniProt (U; see Fig.S1) or RefSeq GFF3 (R): **human MACF1**: MACF1a1 (GUR), MACF1a2 (none/different N-term), MACF1a3 (GR), MACF1b (none/different N-term GUR), MACF1c (R), MACF1-4 (none); **mouse MACF1**: MACF1a1 (GUR), MACF1a2 (G), MACF1a3 (GUR), MACF1b (none/different N-term GUR), MACF1c (partial G), MACF1-4 (none); **human DST**: DSTa1 (GR), DSTa2 (R), DSTa3 (R), DSTb1 (GUR), DSTb2 (R),DSTb3 (R), DSTe (G, starting within Plakin), DSTe2 (R); **mouse DST**: DSTa1 (G), DSTa2 (R), DSTa3 (none), DSTb1 (GU), DSTb2 (R), DSTb3 (R), DSTe (GR, starting within Plakin). Note that all databases annotate additional isoforms which differ from the models proposed in the current literature.

**Fig. S3.**
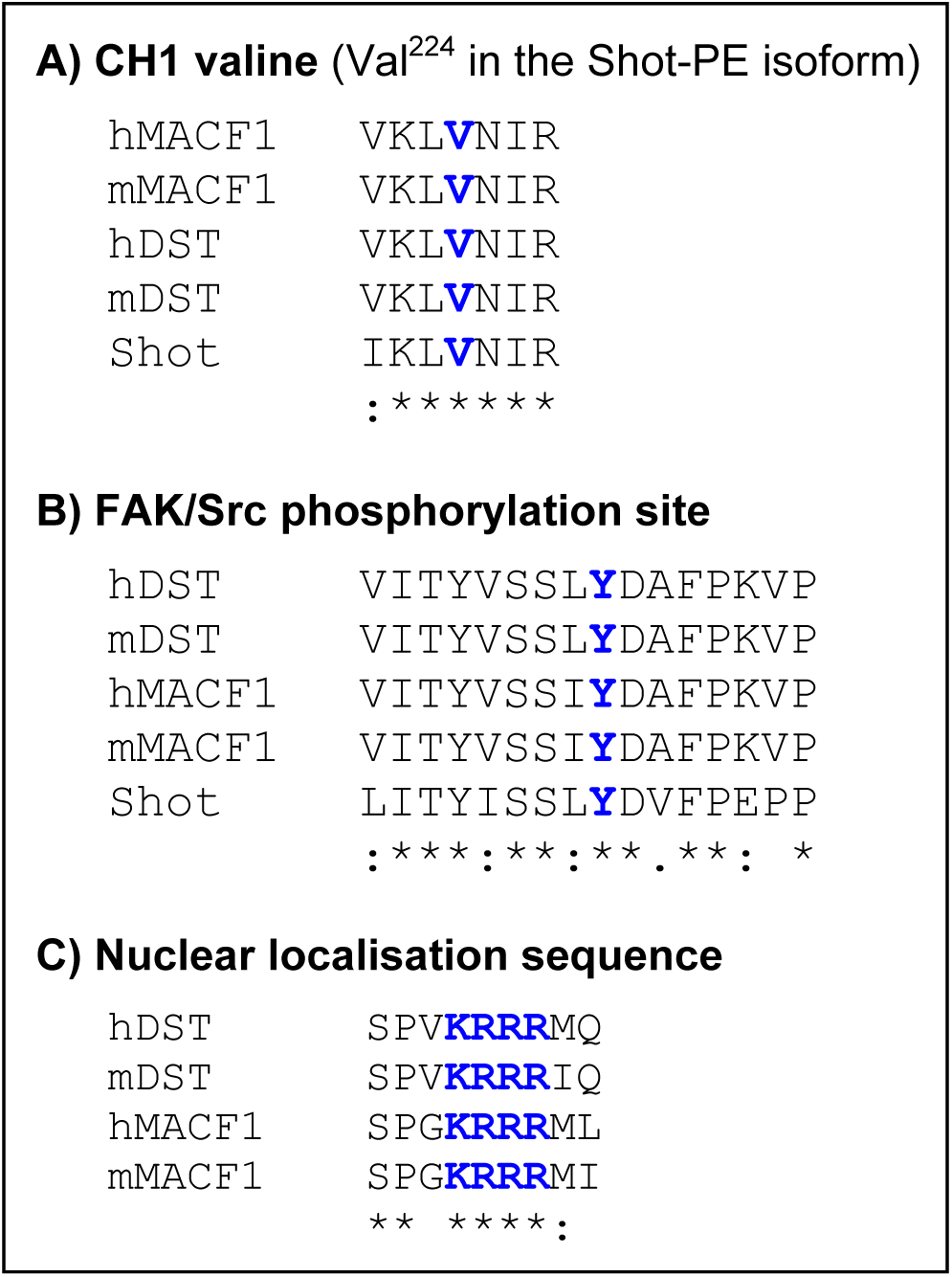
Conserved N-terminal motifs of fly and mammalian spectraplakins. Short genomic sequences are shown for *Drosophila* Shot and human/mouse (h/m) MACF1 and dystonin (DST). **A)** Valine^224^ (bold, blue V) in the first calponin homology domain of Shot’s abolishes binding of the ABD to F-actin [3]; this valine and its surrounding sequence are well conserved in mammalian spectraplakins. **B)** A target site for FAK phosphorylation was first reported for mMACF1 (bold, blue Y is the phosphorylated residue) [4]; it appears evolutionarily well conserved. **C)** A nuclear localisation sequence (NLS; bold, blue) was first reported for the PD of mDST [5] and seems conserved in MACF1 but not Shot (although different NLS are predicted also for the Shot PD). Symbols used: asterisks, identical residues; colon, conservative amino acid exchange; dot, semi-conserved amino acid exchange.

**Fig. S4.**
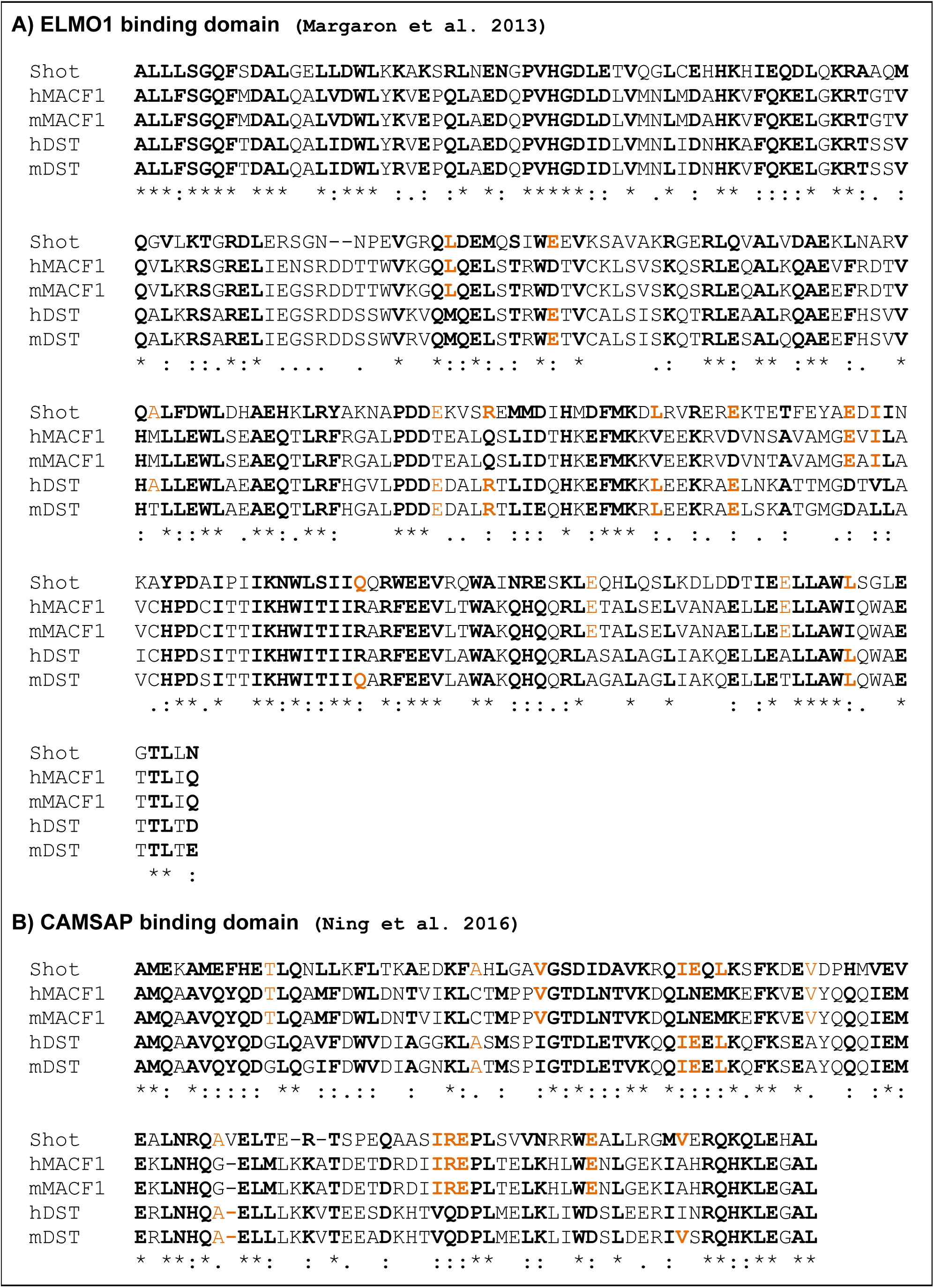
Specific conserved sequences in the SSR of fly and mammalian spectraplakins. Genomic sequences are shown for *Drosophila* Shot and human/mouse (h/m) MACF1 and dystonin (DST). **A)** A region with binding capability for ELMO1 was reported for the SRR of mMACF1 [6] (sequence shown here deduced from provided primer information); it appears well conserved in mammalian and fly spectraplakins (39/57% similarity/identity between fly and mammalian sequences), suggesting that dystonin may interact with ELMO family proteins and Shot with the ELMO homologue Ced-12. **B)** A region with binding capability for CAMSAP was reported for the SRR of hMACF1 [7] (sequence corresponds to ACF7-IV-3(2) 4023-4134; Fig.S3 therein); it appears well conserved in mammalian and fly spectraplakins (35/55% similarity/identity between fly and mammalian sequences). Symbols used: asterisks, identical residues; colon, conservative amino acid exchange; dot, semi-conserved exchanges; orange, amino acids that are identical to only a subset of Spectraplakins.

**Fig. S5.**
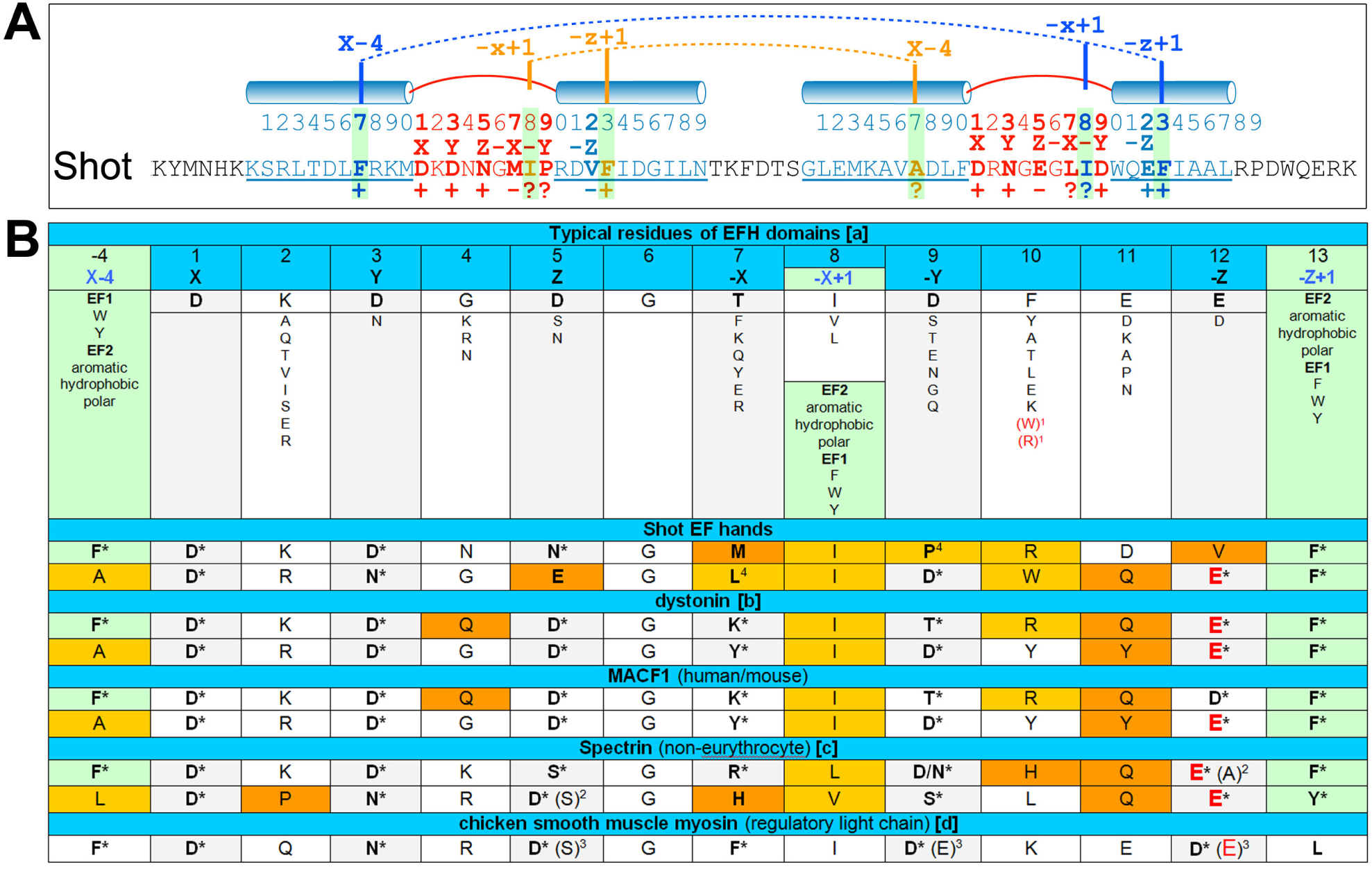
Potential Ca^2+^ binding properties of Shot’s EFH. **A**) A linear model of a paired EFH with two EF motifs, each formed by an ingoing and outgoing α-helix (light blue) connected by a loop (red); the residues at positions 1 (X), 3 (Y), 5 (Z), 7 (-X), 9 (-Y) in the loop and position 12 of the outgoing helix (-Z) coordinate with Ca^2^+; combinations of the residues X-4, -X+1 and -Z+1 connect into cluster 1 (blue) and cluster 2 (orange) and promote antiparallel organisation and hydrophobic packing favourable for Ca^2^+ binding [8, 9]. The shown Shot sequence (position 4763-4808 of isoform E) is predicted to have α-helices in the correct positions (underlined blue; as predicted by 3D-PSSM) and displays many of the typical/required residues (+, in agreement;?, in potential agreement as explained in B; -, not in agreement). **B**) At the top is a list of typical residues (see references in [a] below) and sequences of a number of EFHs are listed below as indicated; symbols with asterisk meet the criteria, fields in dark orange appear not to meet any criteria, fields in light orange are likely to meet criteria given for X-4, -X+1 or -Z+1 [8, 9] or for other residues (as indicated by specific notes explained below); the red, enlarged E in position 12 may be especially favourable for Ca^2^+ binding (see note 3 below). **References**: [**a**] for 1-12 [10], for X-4, -X+1 and -Z+1 (light green fields) [89]; [**b**] [11]; [**c**] [12, 13]; [**d**] [14]. **Notes**: (**1**) valid residues according to Prosite prediction algorithm; (**2**) in the EFH of spectrin, E12A (1^st^ motif; E33A) has a strong and D5S (2^nd^ motif; D69S) a weak impact on Ca^2^+ affinity [13]; (**3**) in the sole EFH motif of chicken smooth muscle myosin, D5S and D9E strongly reduce Ca^2^+ affinitiy, whereas D12E enhances it above wildtype levels, even when in double- or triple-mutant conditions with D5S and/or D9E [14]; (**4**) L at -Y and P at -X are similarly found in calmodulin [9].

**Fig. S6.**
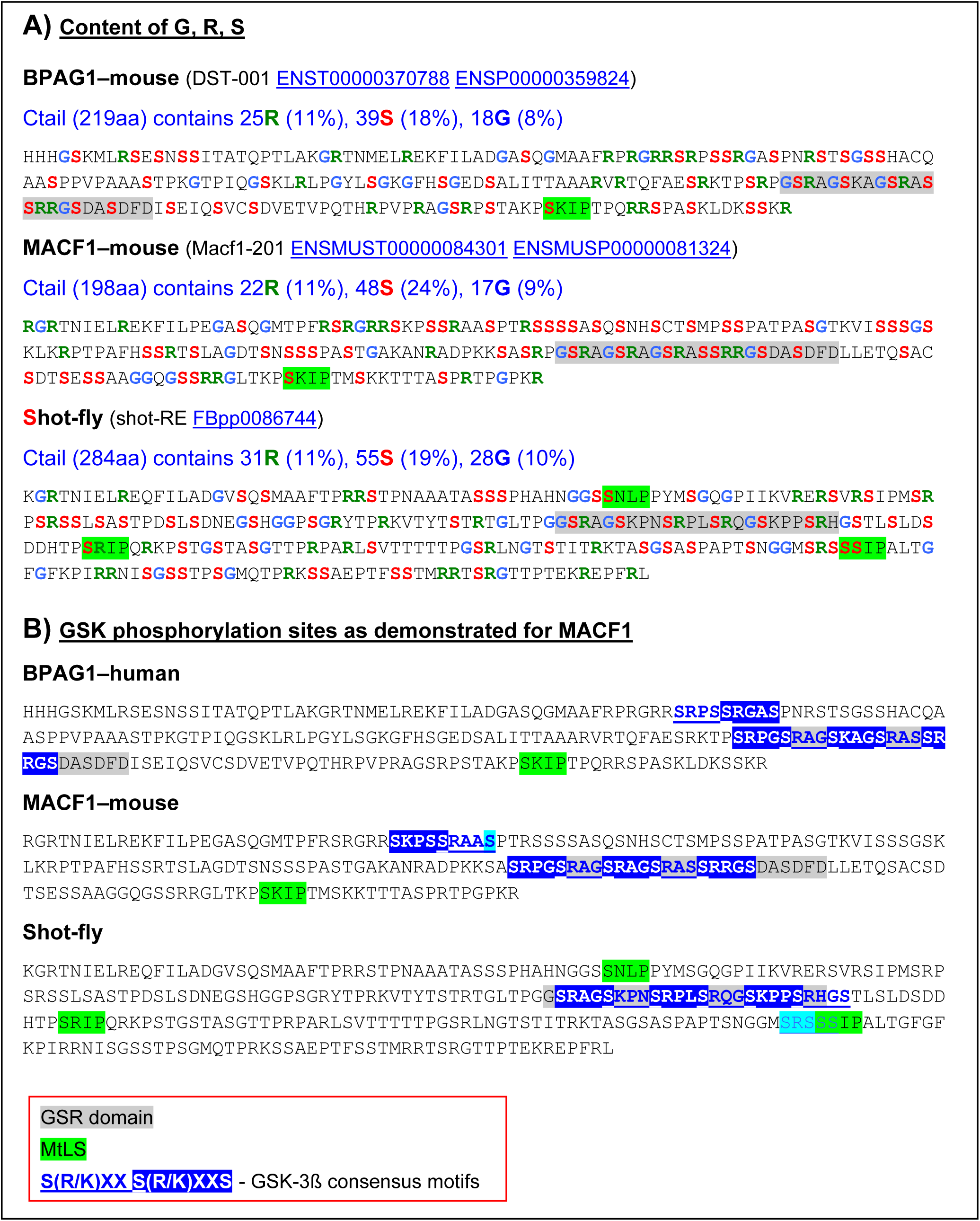
Properties of spectraplakin Ctails. **A**) Even distribution of arginins (R), serines (S) and glycins (G), the position of the GSR domains (grey) as originally predicted [15], and of MtLS (green) [16]. **B**) The GSR domains (grey) overlap with GSK-3ß phosphorylation target sites (blue) identifiable in all three spectraplakins (details in the main text).

